# Planar differential growth rates determine the position of folds in complex epithelia

**DOI:** 10.1101/515528

**Authors:** Melda Tozluo◻lu, Maria Duda, Natalie J. Kirkland, Ricardo Barrientos, Jemima J. Burden, José J. Muñoz, Yanlan Mao

**Affiliations:** UCL/MRC Laboratory for Molecular Cell Biology, University College London, Gower Street, London WC1E 6BT, United Kingdom; Mathematical and Computational Modeling (LaCàN), Universitat Politècnica de Catalunya, Barcelona, Spain; Institute for the Physics of Living Systems, University College London, Gower Street, London WC1E 6BT, UK

**Keywords:** computational modelling, tissue mechanics, finite element, folding, morphogenesis

## Abstract

Folding is a fundamental process shaping epithelial sheets into 3D architectures of organs. Initial positioning of folds is the foundation for the emergence of correct tissue morphology. Mechanisms forming individual folds have been studied, yet the precise positioning of the folds in complex, multi-folded epithelia is an open question. We present a model of morphogenesis, encompassing local differential growth, and tissue mechanics to investigate tissue fold positioning. We use *Drosophila melanogaster* wing imaginal disc as our model system, and show that there is spatial and temporal heterogeneity in its planar growth rates. This planar differential growth is the main driver for positioning the folds. Increased stiffness of the apical layer and confinement by the basement membrane drive fold formation. These influence fold positions to a lesser degree. The model successfully predicts the emergent morphology of *wingless spade* mutant *in vivo*, via perturbations solely on planar differential growth rates *in silico*.

## Introduction

Epithelial folding is a fundamental morphological process that is encountered abundantly during the development of multiple organisms. It is used to sculpt organs from flat epithelial sheets into complex structures such as tubular, undulated, and branched tissues (Nelson, 2016). Folds may function as a means of compartmentalisation, surface area increase to facilitate material exchange, or may emerge as a side effect of pathology, such as overgrowth in cancer (Gutzman et al., 2008; Hruban et al., 2000; Nelson, 2016).

Possibly the most extensively studied driver of folding is apical constriction via accumulation of non-muscle myosin II (Dawes-Hoang, 2005; Granholm and Baker, 1970; Lecuit and Lenne, 2007; Lewis, 1947; Polyakov et al., 2014). Basal relaxation, lateral constriction (Štorgel et al., 2016; Sui et al., 2012, 2018; Wang et al., 2016; Wen et al., 2017) and cell shortening (Conte et al., 2012; Gutzman et al., 2008; Sherrard et al., 2010), constriction of the regions surrounding the prospective fold (Kondo and Hayashi, 2013; Röper, 2012), and constriction of supporting structures by other cells (Hughes et al., 2018) have all been demonstrated as potential folding mechanisms. Other force generation mechanisms, such as cell rounding in mitosis, adhesion shifts, or basal extrusions can also induce folds (Kondo and Hayashi, 2013, 2015; Wang et al., 2012). In all these scenarios, what defines the position of the prospective fold is a biochemical signalling mechanism responsible for selecting the cell population to actively generate the forces.

Beyond cellular forces, confinement by differential growth between different layers of cells can induce instabilities to generate patterns of buckling. This can be observed in a multitude of systems, such as the folding of the gut (Savin et al., 2011), dental epithelium (Marin-Riera et al., 2018), brain (Tallinen et al., 2014), lungs (Kim et al., 2015), and the rippled edges of plant leaves (Dervaux and Ben Amar, 2011; Liang and Mahadevan, 2009; Marder et al., 2003). External structures, such as the extracellular matrix (ECM) surrounding the tissue, can also provide sufficient confinement to growing tissue to induce folding (Diaz-de-la-Loza et al., 2018; Sui et al., 2012). Uniform growth and constriction will induce folds in predictable patterns following the physical rules of buckling (Karzbrun et al., 2018; Pocivavsek et al., 2008; Shyer et al., 2013; Wang and Zhao, 2015). The patterns then can be refined by further perturbations such as local constriction by smooth muscles (Kim et al., 2015), local ECM alterations, adhesive forces, or the overall shape of the tissue (Tallinen et al., 2016). Consequently, dynamic modifications of the ECM and basement membrane (BM) are utilised in large scale tissue morphogenesis, including folding (Diaz-de-la-Loza et al., 2018; Sui et al., 2012).

While shaping a tissue, various mechanisms are likely to occur in parallel, such that once a fold is initiated in a selected position, a combination of modifications of the confinement, active force generation, and cell shape changes can help its progression. A key question is how are the initial positions of the folds defined to achieve the precise tissue morphology (Nelson, 2016)?

As an emergent mechanical phenomenon, fold position selection is likely to depend on a combination of the forces accumulating in the growing tissue, the dynamics of surrounding structures, and the inherent properties of the tissue such as stiffness or shape prior to folding. None of these factors are trivial to investigate independently in an experimental system – how would one eliminate the influence of the shape of a tissue on its form? Therefore, the topology of folding morphogenesis is a problem particularly suitable for computational exploration.

*Drosophila melanogaster* is an established model system for studying morphogenesis. The wing imaginal disc of *Drosophila* forms three distinct folds, perpendicular to the dorsal-ventral axis. These major folds are highly reproducible in their number and positions, marking the boundaries between the notum, hinge, and pouch regions of the wing disc (Fig. 1). There is evidence that basal relaxation, lateral constriction, and stiffness changes within the cell compartments play roles in generation of the folds (Sui et al., 2012, 2018; Wang et al., 2016). However, what determines their positions, and drives the initiation of these folds is an open question. This makes the wing disc an ideal experimental system to investigate general mechanisms that control the position of folds in complex epithelia, a problem that has been under-investigated, but critical in determining the final functional architecture of the tissue.

**Figure 1.**
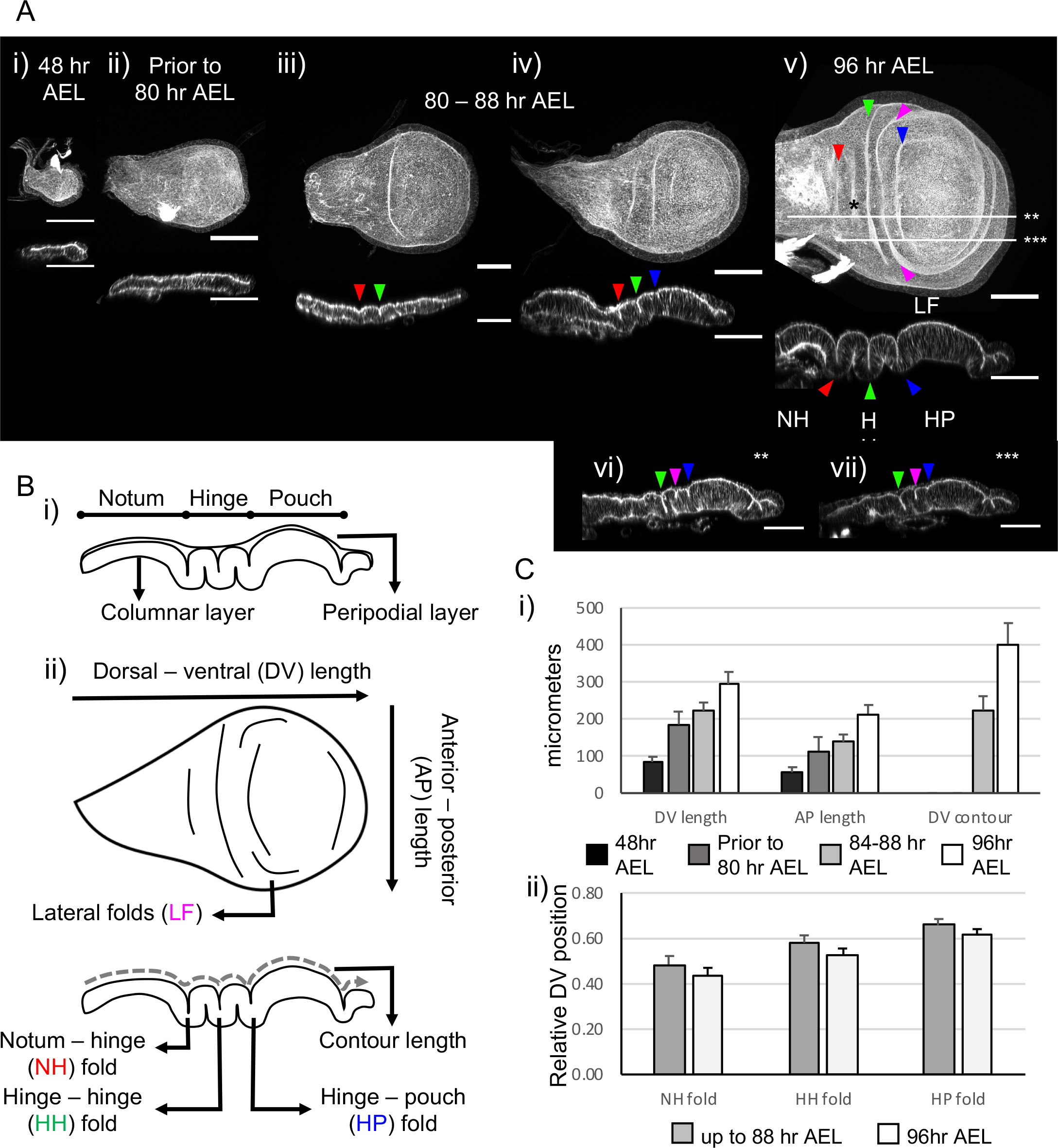
Characterisation of wing imaginal disc fold morphology. A) The morphology changes of the wing imaginal disc between 48 and 96 hours AEL. Top panels: top view, bottom panels: sagittal view. Maximum projections images, actin labelled with fluorescent phalloidin. Arrowheads that point to HN, HH, HP folds are red, green and blue respectively. Lateral folds in between the PH and HH folds are marked with magenta arrowheads. Scale bars are 50 micrometres. i) 48 hr AEL, ii) prior to fold initiation, up to approximately 80 hr AEL, iii-iv) time range between 80-88 hr AEL when folds start to initiate. ii) Representative image with HH fold as an established apical indentation, NH and PH folds starting to initiate. iv) Representative image with all three folds in form of apical indentations, arrowheads same as iii. v) 96 hr AEL, fold morphology complete. Due to the projection, basal folds are visible on the top view, example marked by black star on (v). vi&vii) Lateral cross-sections along lines marked with stars on (v) demonstrating the HH, HP folds, and the LF, vi) **, vii) ***. B) Schematic representing the wing disc structure. i) Domains of the wing disc, the thin peripodial layer is hardly visible on the experimental images. The columnar layer is modelled in our simulations. ii) Folded wing disc, top and sagittal views, with developmental axes, and fold names labelled. C) i) Wing disc size during fold formation, developmental age goes from black to white, see methods for n. At 48 hr AEL, the AP and DV lengths are 56 and 84 μm, respectively. Prior to approximately 80 hr AEL, the lengths are 114 and 185 μm; at 88 hr AEL they become 128 and 222 μm. At 96 hr AEL, AP and DV lengths are 214 and 294 μm, respectively, while the apical contour length on the DV axis is 402 μm. ii) Positions of the folds normalized to dorsoventral (DV) length, error bars represent one standard deviation. Up to 88 hr AEL, the positions are 0.48, 0.58, and 0.66 for NH, HH, and HP folds respectively. Similarly, the positions are 0.43, 0.52 and 0.61 for 96 hr AEL. For 72-88 hours AEL, NH fold n=22, HH fold n = 29, HP fold n=18; for 96 hours AEL, n=14 for all folds.

Here we investigate the minimum set of requirements for initiating the correct topology of the complex multi-folded wing disc epithelia. Our search allows us to postulate planar differential growth as a novel mechanism for fold initiation. We measure the differential planar growth rates through late second instar and early third instar of wing imaginal disc development (48 to 96 hours after egg laying, AEL) with high spatial resolution. Utilising a computational approach, and from experimental measurements, we demonstrate that the differential growth in the plane of the tissue, under the compression of the extracellular matrix, drives initiation of three folds from the apical surface. The differential growth in the height, specifically the relative thickening of the pouch region, helps to constrain the folds to the hinge. We predict that a reduction of early growth in the hinge region, prior to any folds being visible, can affect the number and position of folds that form later. We experimentally validate this prediction against a *wingless* mutant, which has reduced proliferation specifically in the hinge region (Neumann and Cohen, 1996). Our simulations show that the alterations of planar growth rates are sufficient to explain the observed fold perturbations, namely loss of one fold, of the wingless mutant.

## Results

### Characterisation of wing imaginal disc fold morphology

The wing imaginal disc is a monolayered epithelial sac that wraps around itself, forming effectively two layers of cells. The peripodial layer, positioned as the top layer throughout this paper, is formed of squamous cells. The bottom layer is the columnar layer, which forms the folds by the end of third instar (Fig. 1Aiv-v, B). The apical surfaces of both layers face each other towards the lumen, and the basal surfaces face outwards. Both apical and basal surfaces harbour ECM of different compositions (Pastor-Pareja and Xu, 2011; Ray et al., 2015).

To analyse the stages of morphogenesis of the wing disc we focus our attention on the columnar layer and characterise the two-day period from 48 hours AEL, when the tissue is flat and relatively small (Fig. 1Ai), to 96 hours AEL, when the columnar layer has formed three folds at the hinge region, between the wing pouch and the notum (Fig. 1Av, B). From dorsal to ventral tips, these folds are termed notum-hinge (NH) fold, hinge-hinge (HH) fold, and hinge-pouch (HP) fold (Fig. 1Bii). Between the HH and HP folds, tissue forms additional lateral folds (LF) that do not reach to midline (Fig. 5Avi-vii, Bii). There are multiple smaller folds at the ventral tip of the wing disc, where the tissue loops to connect columnar layer to the peripodial layer. These smaller folds are beyond the scope of the current work, as the largely unknown dynamics of the peripodial layer are not included in the current model.

In our analysis, we segment the development into three morphological stages (Fig. 1A, S3A), i) the early stage before initiation of folds, ending prior to approximately 80 hours AEL. During this stage, the tissue is grows relatively flat and folds do not start forming (Fig. 1Aii, Ci). ii) The intermediate stage where the folds are starting to initiate on the apical surface, with the possibility that the HH fold has fully formed, ending by 88 hours AEL (Fig. Aiii-iv, Ci). iii) The stage where all folds are established, ending approximately 96 hours AEL (Fig 1Av-vi). Correlating with the fold formation, DV contour length increases more rapidly than the projected DV length between 88 to 96 hours AEL (Fig. 1Ci). The folds are formed at reproducible, precise positions as normalised to the tissue DV length (Fig. 1Cii).

We match the initial state of our simulations to the tissue size and shape at 48 hours AEL and start simulations using simple growth rates derived from the changes in dimensions of the wing disc as described above (Fig. 1Ci). Step by step, we add in external confinement with apical ECM and BM, physical property heterogeneities, and fine growth patterns to characterise the requirements of fold initiation in the wing disc.

### The computational model

For the purposes of identifying the mechanisms driving wing disc folding, we developed a finite element model of tissue morphogenesis. In our model, the tissue is treated as a non-homogeneous continuous material, and defined as a set of elastic elements that can grow and vary in size from subcellular to multicellular (Fig. 2A-C, S1A). A neo-Hookean material model with time-constant material parameters is selected to represent the stress-strain relationship of the tissue. The cellular and non-cellular (BM) sub-compartments utilise different set of parameters but satisfy the same governing equations. When applied, an external viscous resistance proportional to the exposed surface area and velocity is employed. Discrete form of transient balance equations give rise to a system of non-linear equations that is solved numerically with a Newton-Raphson method. Ranges of elastic and viscous properties have been tested and presented in following sections. For all simulations, the Poisson ratio is taken to be 0.29 (Schluck et al., 2013).

**Figure 2.**
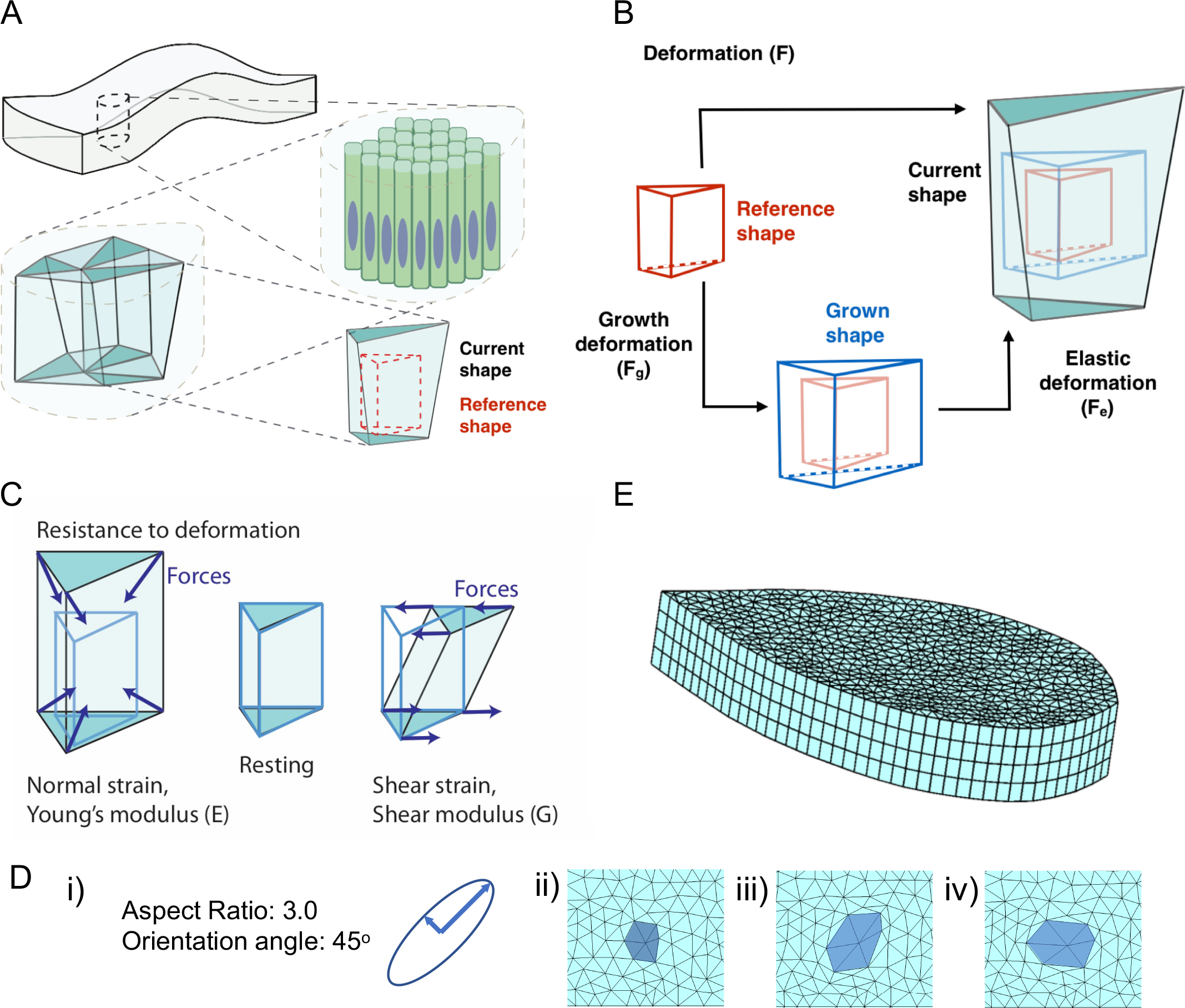
The computational model design. A) Schematic describing the definition of finite elements. Each element represents a continuous material slab of the tissue, the elements are independent of cells, the size can be multicellular or subcellular. B) Schematic of growth methodology in the model, red wire plot: the ideal shape of the element at the beginning of the simulation, blue wire plot: the desired shape of the element obtained from the growth of the reference element C) Schematic representing elastic forces generated by elements upon deformation. D) i) The oriented growth input ii-iv) Simulation with a single clone (highlighted in dark blue) growing surrounded by non-growing tissue. ii) Initial state of a tissue fragment, before growth iii) Tissue in (ii) after application of growth in (i). iv) Tissue in (ii) after growing with the same rates at an orientation angle of zero degrees. E) Initial mesh of the simulation, at 48 hour AEL. The mesh is generated with the contour of a sample 48 hour AEL old wing disc, and scaled to tissue dimensions given in Figure 1C. See also Supplementary Figure 1.

A key concept in modelling morphogenesis is the definition of growth. The simulations can utilise spatially and temporally heterogeneous growth rates and growth orientations. Growth is applied on the initial reference orientation of each element, whose total deformation gradient is split onto a growth and an elastic contribution through a multiplicative decomposition (Rodriguez et al., 1994; Taber, 1995) (Fig. 2B). We extended the existing approach to include oriented growth that follows the plane of the tissue. The rotations defining the oriented growth are constructed such that any rotation around the apical-basal axis will be incorporated in the orientation of the growth, ensuring the growth continues in the desired orientation in the plane of the tissue, rather than in fixed spatial coordinates. In contrast, any mismatch between the world (spatial) and Lagrangian apical-basal axis is ignored, as we wish to apply planar growth in the original tissue plane (Fig. 2D, S1B) (See Methods: The model definition).

Remodelling is calculated at element level, where the desired geometry of the element is slowly updated to match its current geometry, relaxing strains in the process. The principal values of oriented growth are utilised to implement remodelling, where the “growth rate” associated with remodelling is defined by the current deformation of the element, and the remodelling half-life.

The initial geometry is defined by tessellation of the contour of a wing disc at 48 hours AEL, scaled to the average dimensions (Fig. 1C, 2E). We assume anterior-posterior symmetry and model half of the tissue. As such, the simulations are run with fixed y-position (along AP axis) for all nodes at the dorsal-ventral plane of symmetry that is at the midline of the tissue. A hard-wall potential is added to the elastic potential for volume exclusion. Further, self-contact is modelled in such a manner that nodes belonging to separate elements, but within a threshold distance of each other can adhere. The nodes within the same element that are within a very small distance of each other are merged, collapsing the edge of the element, to avoid element flipping (Fig. S1C-E). The circumference of the tissue has boundary conditions limiting bending, such that all nodes on the same column at the boundary have the same x & y coordinates (Fig. S1F). Further details or the model and formal definitions are provided in the methods section.

### Resistances from apical basal surface confinement are essential for folding

We start our simulations with uniform growth obtained from the tissue size measurements presented in Figure 1Ci. Here, we calculate constant growth rates that would bring the tissue AP and DV contour lengths from their sizes at 48 hours AEL to 96 hours AEL, in a 48-hour time window (0.028 hr^−1^ in AP, and 0.033 hr^−1^in DV). As expected, our simulations with uniform growth on a tissue with minimal external resistance and homogenous physical properties do not form any folds (Fig. 3A, top-left-corner). This result reinforces that the tissue must have external factors driving compression.

**Figure 3.**
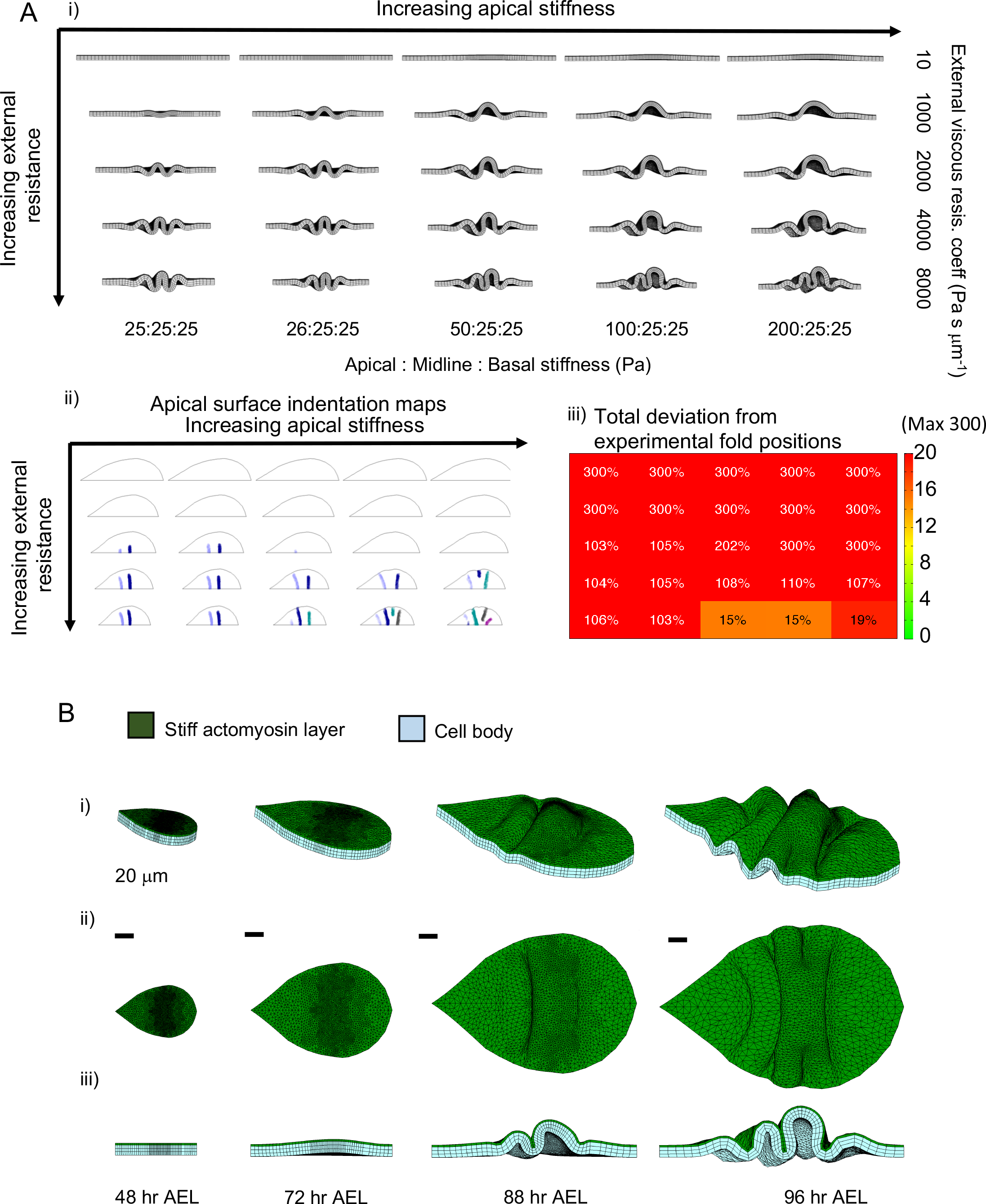
Relative increase in apical stiffness and external resistance to tissue growth are essential for fold formation. A) i) Simulation results for a tissue growing from 48 hour AEL to 96 hours AEL, with uniform planar growth rates. The growth rate is 0.033 hr^−1^ in DV, and 0.028 hr^−1^ in AP, as calculated from Figure 1C. Each image shows the midline of the tissue central line at 96 hours AEL, dorsal tip on the right. Columns: increasing external viscous resistance; rows: increasing apical stiffness relative to the rest of the tissue. Nonsymmetric folding can only occur with relatively high apical stiffness and high external resistance (bottom right corner). ii) Apical indentation maps automatically identified from the curvature of facing surfaces, each continuous folding region is marked in a single colour. iii) Fold position deviations, calculated as sum of percentage deviation from each experimental fold at the tissue centre. Missing folds count as 100 per cent deviation, maximum deviation being 300%. The colourbar scans a lower deviation range, up to 20 per cent. For ii and iii, the grid organisation is same as in (i) B) Simulation with apical stiffness 200 Pa and external viscous resistance coefficient applied to both surfaces at 8000 Pa s μm^−1^. i) Orthogonal perspective view, ii) top view, sagittal view as cross-section at midline. Scale bars are 20 micrometres. Timeline as depicted under each the panel. See Movie 1, and Supplementary Figure 2.

During morphogenesis, both apical ECM and BM can exert resistances to tissue growth and movement, and this resistance is functional in the formation of correct tissue architecture (Diaz-de-la-Loza et al., 2018; Hannezo et al., 2015). Initially, we represent this as a viscous external resistance on the apical and basal surfaces of the growing wing disc. As the viscous resistance is increased, the tissue starts forming ripples with compact, regular folds, initiating from the centre and distributing towards the dorsal and ventral edges (Fig. 3Ai, first column). To assess the geometry of the fold formation, we generate apical fold initiation maps. Here, fold initiation is automatically detected by the curvature of the facing surfaces, and the identified indentations are marked on the tissue outline with each continuous indentation given in a single colour (Fig. 3Aii). Then the total deviation in fold positions is calculated as percentage of the DV length, each missing fold deviation contributing 100 per cent to the sum (Fig 3Aiii). Within the tested range with uniform tissue physical properties (Fig. 3Ai-iii, first columns), the tissue can form only two folds. The amplitude of the folds brings their peaks higher than the apical surface of the pouch and notum regions, which is not the case for the wing disc, where the hinge folds form within the thickness range of the notum and pouch.

Next, we considered physical property heterogeneities within the wing disc as a source of breaking symmetry and improving emergent number and morphologies of folds. It is likely the imaginal disc epithelium has heterogeneities, especially along its apical-basal axis. The dense actomyosin mesh on the apical surface, or the actin of the basal surface could both potentially have higher stiffness than the rest of the cell body (Farhadifar et al., 2007; Sui et al., 2018). Alternatively, accumulation of the cell nuclei in the middle zone of the wing disc could effectively bring about a stiffer tissue midline(Meyer et al., 2011). To account for all these possibilities, we simulated wing disc growth with an increased stiffness on the apical surface (Fig. 3A), on apical and basal surfaces (Fig. S2A), or on the midline (Fig. S2B). Of the tested cases, only stiffness increase on the apical surface could generate three folds; with 15-19 per cent deviation from the correct fold positions at tissue midline (Fig. 3Ai-iii bottom rows, B, Movie 1). However, none of the physical property heterogeneities were sufficient to generate folds similar to the experimental morphology, which has three folds on the hinge that have amplitudes comparable to the rest of the tissue thickness (Fig. 1Av).

With this analysis, we conclude that external resistance to growth is essential for buckling the tissue and that increased apical stiffness can induce correct number of folds. However, defining growth rates as uniform and the BM as a simple viscous resistance is not sufficient to induce folds in the correct positions and shapes. Therefore, we constructed detailed maps of tissue growth at fine spatial and temporal resolutions to improve the implemented growth rates.

### Wing imaginal disc planar growth patterns show spatial and temporal heterogeneity

With the purpose of improving the definition of growth in our simulations, we experimentally measure the local growth rates via clonal analysis. By inducing sufficiently sparse single cell clones at different ages AEL, and dissecting the tissue 24 hours later, we quantify the local growth rates and generate spatial growth maps (Fig. 4A). For each clone, the extent of growth is defined by the number of nuclei in the clone; the orientation and aspect ratio of growth are defined by the ellipse fitted to the clone shape (Fig. 4Aii). The centre of the fitted ellipse defines the position of the measured growth as normalised to the tissue bounding-box. Assuming symmetry in the anterior-posterior axis, all single clone measurements are binned on a 2D grid defined on half of the tissue bounding box (Fig. 4Aiii), which is then reflected to the remaining half. This analysis resulted in coupled growth rate heatmaps and orientation maps, defining the growth patterns of the tissue in fine spatial detail (Fig. 4B). We have repeated this process in three time windows as identified from the morphological quantifications (Fig. S3A). To improve alignment of measurements from different experiments and improve spatial accuracy, we aligned each experiment to the HH fold at stages when this fold is identifiable (i.e. mid and late growth phases) (Fig. S3B). The growth rates of the pouch region of the wing disc have previously been characterised in high detail by Mao *et. al.*(Mao et al., 2013). Measuring the position of the wing pouch at each stage, we overlaid these measurements onto our growth and orientation maps (Fig. S3C), resulting in our final spatial-temporal growth profiles to be utilised in our simulations (Fig. 4B, S3Aiii/ E).

**Figure 4.**
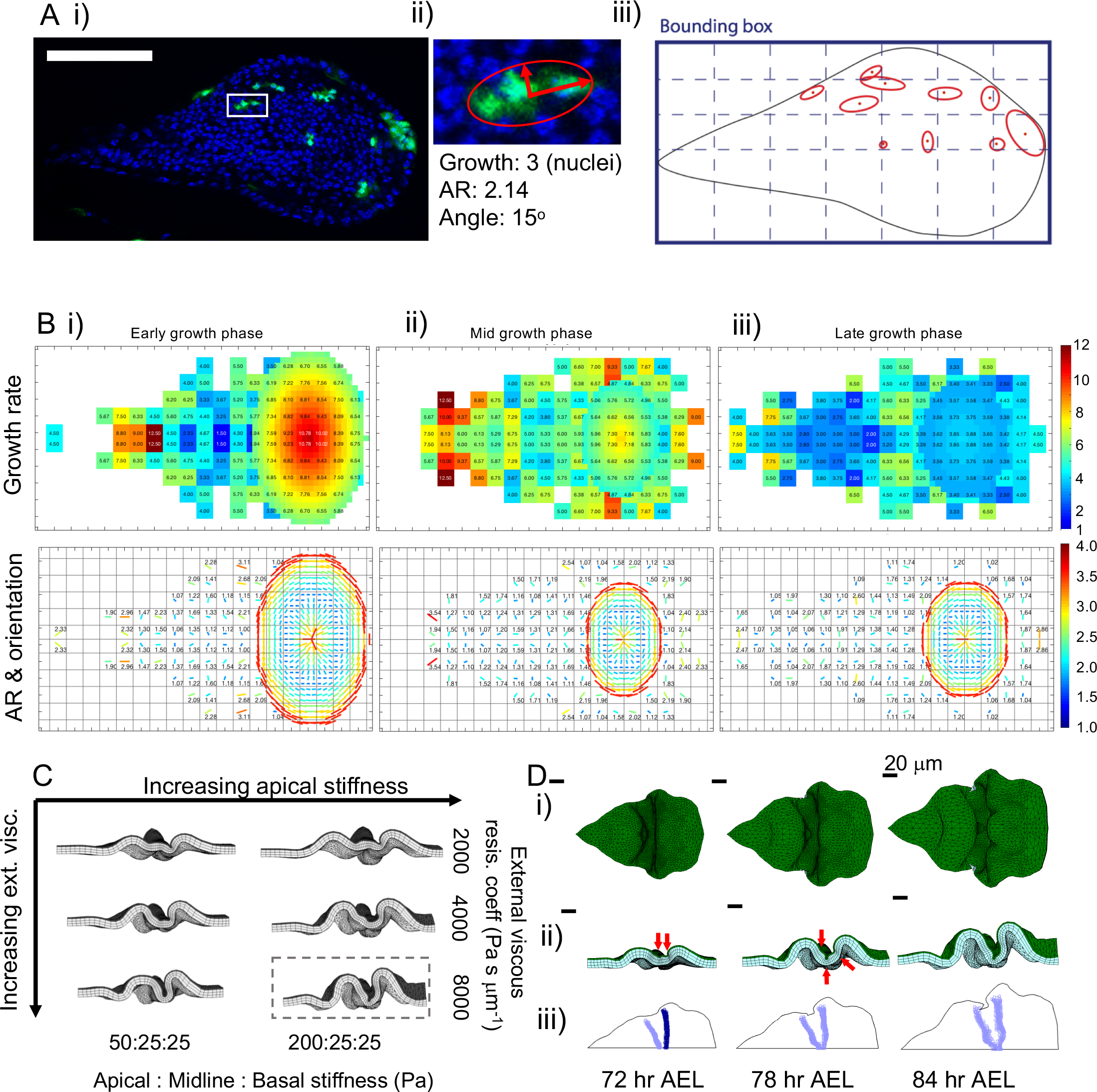
Wing imaginal disc planar growth patterns show spatial and temporal heterogeneity. A) i) Wing disc tissue with nuclei labelled with DAPI (blue), sparse single cell clones induced at 48 hours AEL (green), and disc dissected at 72 hours AEL. ii) Close up of clone marked by the white box in (i), fitted ellipse (red) orientation and aspect ratio noted. iii) Schematic showing fitted ellipses of all clones in (i), mapped on to a grid on the bounding box of the projected image. Contour of the tissue marked within the bounding box. Symmetry in anterior-posterior axis is assumed. B) Growth maps for overlapping time ranges, aligned for fold positions, and with pouch growth data from Mao *et al*, 2013 added. Top panels, growth rate defined by the average number of nuclei in each grid point. Bottom panels, growth orientation maps. The major axis of the average fitted ellipse are shown with lines. The length and colour of the line represent the aspect ratio, orientation of the line represents growth orientation angle. Colourbars for each row are on the right-hand side. i) 48-80 hours AEL, ii) 56 – 88 hours AEL, iii) 72-96 hours AEL. C) Simulations with experimental growth rates and apical and basal confinement modelled as viscous resistances. Increasing apical stiffness on rows, and increasing external viscous resistance coefficient in columns. D) Simulation snapshots for the boxed parameter set in (C). i) top view, ii) Sagittal view as cross-section from the DV axis midline. Indentations starting to emerge (red arrows), the overall morphology does not resemble the experimentally observed pattern. iii) Folded regions automatically identified from the curvature of facing surfaces, each continuous folding region is marked in a single colour. See Movie 2 and Supplementary Figure 3.

The analysis revealed that the wing disc harbours high spatial and temporal variability in its planar growth patterns. Specifically, notum and pouch regions have significantly higher growth than the hinge region at early stages (Fig. 4B, S3E). Similar to the previous observations for the pouch region(Mao et al., 2013), the overall growth rates of the tissue are reduced as the tissue ages.

Next, we implemented these measured growth rates in our simulations, and modelled the 48-hour development (48 – 96 hours AEL) where the measured planar growth rates are applied sequentially in three equal time windows (Fig. S3Aiii). With apical and basal confinement from viscous resistances, the experimental growth patterns result in folding patterns distinct from those of the uniform growth (Fig. 4C-D, Movie 2). Some indications of apical indentations emerge around the hinge region (Fig. 4Dii, red arrows), and the tissue generates two lateral apical indentations that merge at tissue midline in later stages (Fig. 4Diii). None of tested cases can initiate three distinct folds. This suggests that additional to the spatial-temporal variability in growth patterns, the BM should also be modelled in finer detail.

### Characterisation and explicit definition of the basement membrane of wing discs

To characterise the morphology of the BM structure, we acquired electron microscopy (EM) images of wing discs at pre-folding stages (72 hr AEL) and at the end of third instar (120 hours AEL) (Fig. 5Ai-ii) and quantified the thickness of the BM (Fig. 5Aiii). The images reveal the BM is an approximately 0.1μm thick uniform sheet in young discs. For older discs the BM is a more complex structure with multiple layers, seemingly an almost mesh like web, topped with a homogenous thin layer, and a total variable thickness in the range 0.4-0.6μm. Assuming a constant thickening rate of BM at the pouch centre, the average thickness between 72 and 96 hours AEL is then 0.2 μm. Based on these measurements, we defined the BM in our model as a 0.2 μm thick elastic layer encapsulating the tissue (Fig. 5B).

**Figure 5.**
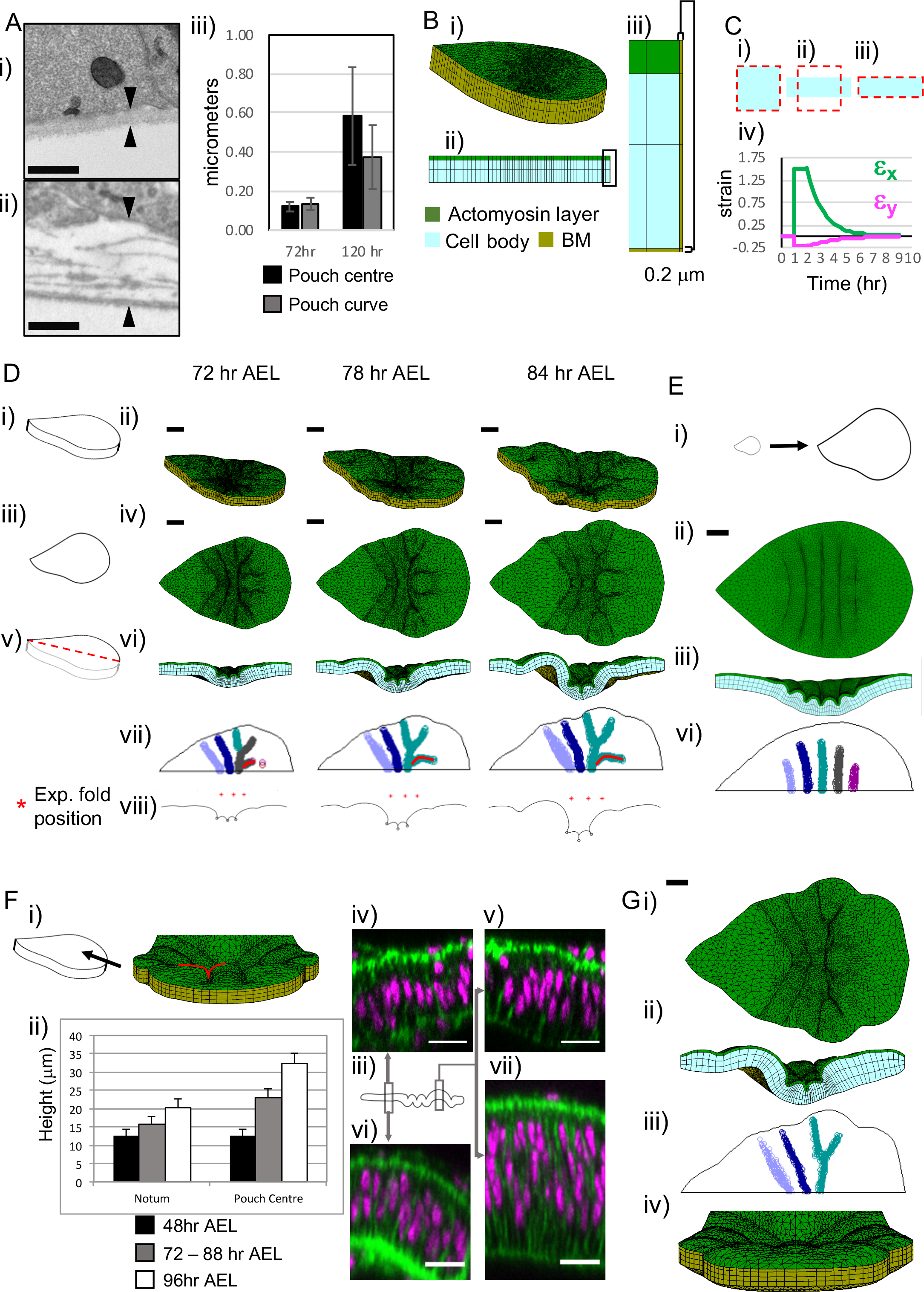
An explicitly defined BM, with planar differential growth rates, enables emergence of *in vivo* mimetic fold morphology. A) Electron microscopy images of the wing disc, i) 72 hours AEL, ii) 120 hour AEL, the regions below the pouch are displayed. Arrowheads mark the thickness measurement. Scale bars are 0.5 micrometre. Also see Figure S5C. iii) The thickness of the BM below the centre (black) and side curve (grey) of the pouch, at 72 and 120 hours AEL, error bars represent one standard deviation. B) Initial simulation mesh with the addition of BM. i) Perspective view, ii) sagittal view, iii) close up of the boxed area in (ii). The actomyosin layer marked in green, rest of the cell body in blue, and BM in yellow. C) Remodelling exemplified with single 2D element, i-iii) blue quadrilateral is the current shape; dashed red lines depict the corresponding reference state including remodelling. i) Element in resting state, ii) immediately after deformation, corners fixed on the x direction. iii) Same element after area conserving remodelling. iv) Green strains in principal axes plotted against time, first principal deformation axis in green (x in the current scenario), second (y) in magenta. Element deformed at t=1 hour without remodelling, remodelling activated at t=2 hours, with half-life of 1 hour. D) Snapshots from simulation with explicit BM definition, at time points 72, 78 and 84 hours AEL. Apical stiffness is 100 Pa, cell body stiffness is 25 Pa, BM stiffness is 1600 Pa with renewal half-life of 8 hour, apical viscous resistance coefficient is 16000 Pa s μm^−1^, and basal is 10 Pa s μm^−1^. See Movie 3. i) Schematic representing the orthogonal perspective view, ii) simulation snapshots. iii) Schematic representing the top view, iv) simulation snapshots. v) Schematic representing the sagittal view at the cross-section of tissue midline on DV axis, vi) simulation snapshots. All snapshots at the same scale, scale bars represent 20 micrometers. vii) Apical indentation maps for each time point, with the tissue apical circumference in black. Each continuous indented region is marked with the same colour. Ectopic pouch fold is marked with red line. viii) The positions of the folds on the tissue cross section apical surface profile, the red stars mark the experimental fold positions measured for 72-88 hours AEL (Fig. 1Ci) E) Simulation results with uniform growth on the tissue plane, simulation parameters same as (D), growth rate same as Figure 3A. i) Schematic to scale, representing uniform growth from 48 to 96 hours AEL. ii-iii) top and sagittal views at 84 hours AEL, respectively. Scale bar 20 micrometres. iv) Apical indentation map at 84 hours AEL. F) Close up view of tissue pouch from dorsal tip showing the ectopic folding. i) Left, schematic representing the view. Right, simulation snapshot at 84 hours AEL. The ectopic fold is marked with a red line, this indentation is marked with red line in apical indentation maps (Dvii) ii) Plot of tissue height of notum and pouch centre, error bars represent one standard deviation (n=4 for initial, n=14 for early, n=17 for late phases). iii) Schematic representing the positions of the zoomed views of nuclei positions, iv-v) The nuclei positions of the notum(iv) and pouch (v) prior to fold formation and vi-vii) positions at notum(vi) and pouch (vii) at 96 hours AEL, showing the increase in tissue height and nuclei pseudostratification. Nuclei labelled with DAPI (magenta), and actin with phalloidin (green). Scale bars 10 micrometres. Measured at 48 hours AEL, prior to fold formation (~80 hours AEL), and 96 hours AEL, the notum thickness is measured as 12.5±2.17, 15.81±2.46, 20.19±2.95 μm, meanwhile the pouch centre thickness is 12.5±2.17, 22.96±1.86 and 32.2±2.26 μm, (mean and one standard deviation). G) Simulation of tissue growth after addition of z-growth, all remaining parameters are same as (D). Snapshots are from 84 hours AEL, scale bar 20 micrometres. i) Top view, ii) sagittal views, iii) fold profiles, note the disappearance of the perpendicular indentation on the pouch compared to (Dvii), vi) view from ventral tip, as comparison to (Fi). See Movie 4, and Supplementary Figures 4 & 5.

In the simulations, the growth of the BM is defined as remodelling upon deformation. Each BM element grows in the orientation of its current deformation, at a rate set by the local remodelling half-life, as detailed under methods. This leads to gradual relaxation of BM deformations, and an emergent, non-homogenous growth of the BM influenced by both the growth rate and shape changes of the cellular layer. For clarity, the relaxation of deformation with such remodelling is represented in a 2D schematic in Figure 5C, where upon change of the current shape of an element (blue square, Fig. 5Ci-ii), the preferred shape gradually changes and aligns with the current shape (red dashed square, Fig 5Cii-iii), relaxing the strains in the process (Fig. 5Civ). Upon refinement of the BM definition, we simulated the development of the wing disc with a series of BM stiffness, remodelling half-life, and apical viscous resistance coefficient parameters.

### Differential planar growth rates of the tissue constrained by an elastic basement membrane drives precise fold initiation

Upon definition of the elastic BM, we initially investigated apical-basal tissue stiffness heterogeneity ranges (Fig. S4A-C), using the experimentally measured growth rates (Fig. 4B). As we increase the apical viscous resistance, the tissue starts forming buckles (Fig. S4A-C). For the cases with increased apical stiffness and increased stiffness on both surfaces, three apical indentations emerge (Fig. S4A/Ci). Of the two scenarios, increased apical stiffness initiates a more dome-like pouch, proportional hinge fold indentations, and lower percentage deviations from the experimental fold positions (Fig. S4A-Cii), better mimicking the *in vivo* fold pattern.

Simulation snapshots for a setup with 100 Pa apical and 25Pa cell body stiffness demonstrate the emerging indentations and their depth as the development progresses, at 72, 78 and 84 hours AEL (Fig. 5Di-vi, Movie 3). In the apical fold initiation maps, we can see the emergence of all three folds, the curved pouch border marked by the HP fold (grey, then cyan), and the emergence of the lateral fold between HP and HH folds (cyan) (Fig. 5Dvii). The folds are concentrated to the central region of the tissue, close to the experimental fold positions (Fig. 5Dviii), with a total deviation of 4 per cent. We predict the initiation of the folds requires the BM to be orders of magnitude stiffer than the cellular layer, which indeed has been inferred to be the case (Keller et al., 2018). As long as the BM is dramatically stiffer than the cellular layer, this phenotype can be produced with a large range of stiffness and remodelling half-life parameter sets for the BM, relative changes in one compensating for the other (Fig. S5A). Simulating the same parameter set as Figure 5D with uniform growth defined in previous sections reveals numerous uniform symmetric ripples on the apical surface (Fig. 5E), rather than the characteristic three-fold architecture. This signifies the importance of planar differential growth in the precise selection of the number and position of the folds.

While reproducing the initiation of three folds of the hinge region, our simulation also initiates an ectopic buckle on the pouch, which is not observed in live tissue (Fig. 5Dvii-part of cyan pouch fold marked in red line, Fi). To decipher what may be driving the resistance of the pouch region to such buckling, we turned our attention to tissue thickness. Our analysis revealed, through the 48 hours of our interest, the wing disc increases its thickness in a non-uniform manner (Fig. 5Fii), with the pouch region becoming relatively thicker than the rest of the tissue. In simulations, the tissue height increases due to compression, the pouch height becoming 17.3μm, and notum height becoming 13.3μm at 84 hr AEL, from the initial uniform height of 12.5μm at 48 hr AEL. This emergent thickness is well below the experimental observations (Fig.5 Fii). Therefore, this thickness increase should be an active growth input in the model.

Implementing this increase in the tissue thickness, the simulations preserve the fold initiation for all three folds of the hinge region, as well as the lateral fold between HH and HP folds, but the pouch surface buckling is prevented (Fig. 5G, S4D, Movie 4). The total deviation from experimental fold positions is 10 per cent. Our simulations predict that within the mechanical context where the hinge folds are initiated, the relative thickness of the pouch region protects it from further buckling that would be otherwise induced by the compactness.

### Early growth pattern is sufficient for correct fold initiation

Our results so far demonstrate the importance of planar differential growth in defining fold positions. The simulations reveal the fold initiation starts by the end of early growth phase (Fig. S4E). To investigate if the early growth rates, and the related force accumulation, are sufficient to initiate the folds, we run simulations where the early growth rates in Figure 4Bi are applied for the first 16 hours (48 to 64 hours AEL) as in the control case, and then the growth is continued with the uniform rates calculated from overall tissue size change. This simulation reveals the early growth rates followed by uniform growth are sufficient for the emergence of *in vivo* mimicking fold morphology (Fig. S6A). When the simulations are run with control growth rates, but the accumulated forces are relaxed prior to emergence of the folds, the emerging morphology is mildly perturbed. The final topology has loose fold formation, and the pouch curve shrinks compared to the control simulations (Fig. S6B-C). These results suggest that the early growth rates measured prior to initiation of the folds, and the corresponding force accumulation, are necessary and sufficient for correct tissue morphology.

### The simulations successfully predict the disrupted fold morphology upon perturbations of planar differential growth patterns in *wingless* mutants

Next, we decided to challenge our simulations with perturbations in tissue growth. Given the sufficiency of early growth rates for patterning the tissue, we needed to select a target gene that already marks the wing disc prior to formation of any folds. The expression of *wingless* (wg) localises in two concentric rings encapsulating the pouch region from early third instar, before the folds start initiating. *spade*^*flag*^ (*spd*^*fg*^) mutations in wg lead to loss of *wg* expression at the inner ring, coinciding with the hinge. Consequently, the mutant has reduced growth in the hinge, and a reduced size in the corresponding region of the adult wing(Neumann and Cohen, 1996; Rodríguez Dd et al., 2002). To predict the wing disc phenotype of this mutation, we run simulations where hinge growth is reduced to the range between 25 to 75 per cent of wild type. At 50 per cent growth reduction (Fig. 6A), the simulated mutant loses the three-fold morphology and displays only two folds at the midline of the disc (Fig. 6B). On the lateral side, a third fold starts emerging, and collapses on the pouch side fold before reaching the midline (Fig. 6Biii). It follows that the *wg* mutant should be able to form the NH fold, while HH and PH folds and the lateral fold will collapse into a single fold at the tissue midline (Movie 5). To test our prediction, we examined *spd*^*fg*^ mutant wing discs (Fig. 6C). The morphology of mutant wing discs at sequential developmental stages clearly demonstrates the emergence of a two-folded morphology (Fig. 6Di-iii) instead of three. Further, investigating the lateral cross-sections of the tissue at 96 hours AEL shows that a third fold initiates at the lateral regions, yet collapses with the pouch side fold before reaching the tissue midline (Fig. 6Div-v), matching with the model predictions (Fig. 6Biii, Movie 5). Simulations where the hinge growth is reduced to 25 and 75 per cent of the wild type levels reveals a dose dependent perturbation of the fold structure (Fig. 6E). At growth rates as low as 25 per cent of wild type, the third fold does not emerge from the lateral sides (Fig. 6Ei), whereas at 75 per cent, three folds form at the midline, albeit being more compact than the wild type (Fig. 6Eii). Then going back to the experiments, we could identify cases where small peaks emerged at the groove of the pouch-side fold at tissue midline (Fig. 6F). This further supports the hypothesis that the HH fold collapses with the HP fold and the lateral fold in the *spd*^*fg*^ mutant. Depending on the level of growth perturbation, the loss of the HH fold can be observed gradually, again matching with the predictions of the simulations.

**Figure 6.**
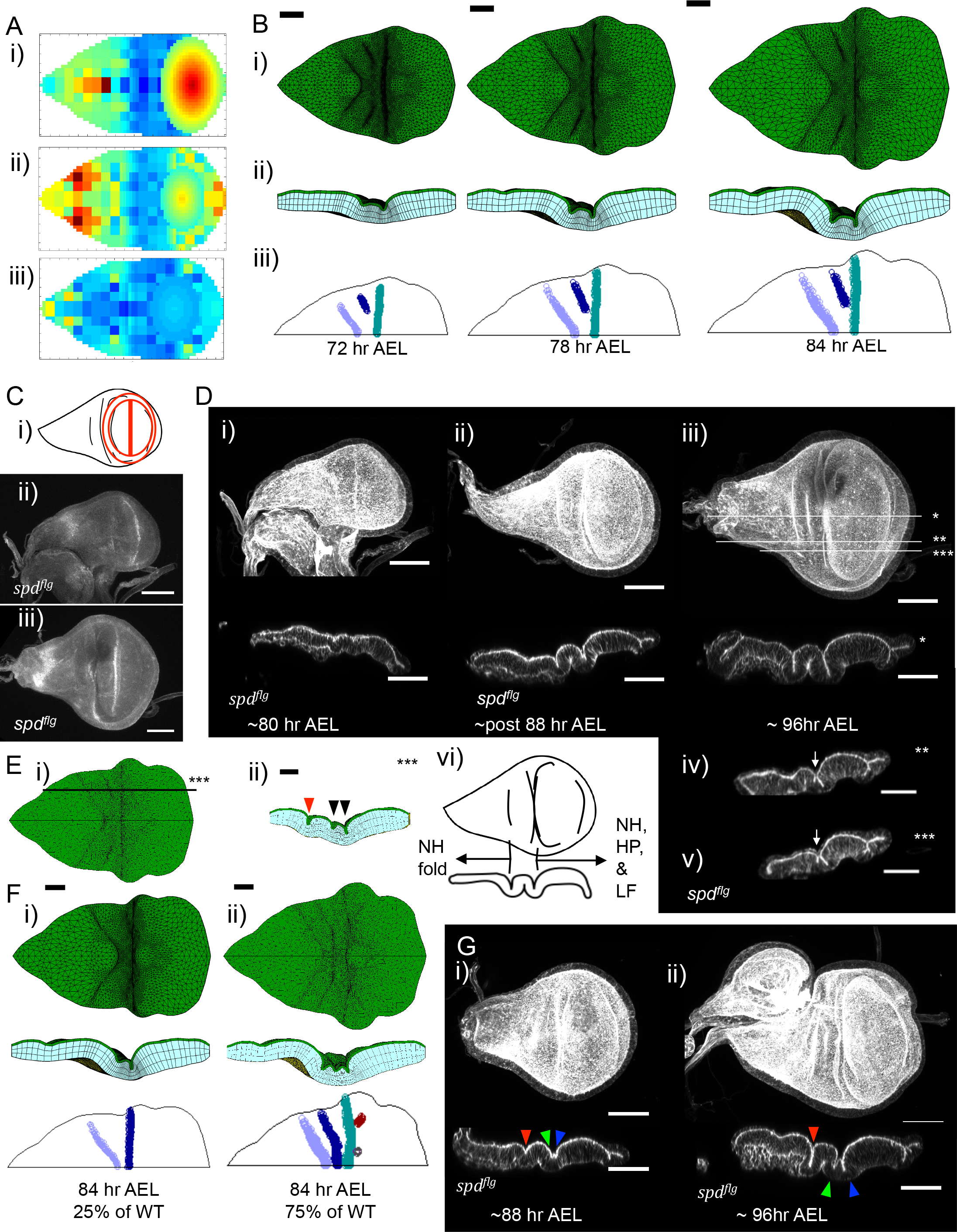
Simulations successfully predict the disrupted fold morphology of *wingless* mutants upon perturbations of planar differential growth patterns. A) Growth maps for *spd*^*flg*^, with 50% reduced growth at the hinge region, defined as a straight band in between the experimental positions of HN fold and centre of the pouch, excluding the pouch itself. The data points sampled from the extended maps (see Figure S4) through the simulation are displayed. Growth measured for i) 48-64 hours AEL, ii) 64-80 hours AEL, iii) 80-96 hours AEL, in line with the utilisation of growth rates given in Fig. S4A. All represented as corresponding growth increase in 24 hours. Colour coding scale same as Fig. 4B. B) Simulation with 50% reduced hinge growth, the maps in (A). i-ii) Simulation snapshots from top (i) and sagittal (ii) view, scale bar 20 micrometres. Simulation times 72, 78 and 84 hours AEL. iii) Apical indentation maps, each continuous indentation marked in single colour, tissue apical circumference in black solid line. All simulation parameters except for the growth rates are same as Fig. 5D. C) i) Schematic marking the pattern of wingless expression in wild type wing discs. The inner ring appears prior to fold formation in early third instar, followed by the outer ring in late third instar. ii) Wingless staining in *spd^flg^*, prior to formation of folds, the stage where the inner ring should have appeared. iii) Wingless staining in late third instar mutant wing disc, showing the outer ring of expression, and lacking the inner ring. Scale bars 50 micrometres. D) The morphology of the mutant wing discs at consecutive ages during the emergence of the folds. Actin labelled with fluorescent phalloidin in grey. Scale bars 50 micrometres. Approximately i) 80 hours AEL, ii) 88 hours AEL, iii-v) 96 hours AEL. iii) Lower panel is the central cross section marked with * in top panel. iv&v) The cross-sections closer to lateral regions, marked as ** and *** on (iii) top panel, respectively. Arrows point to the additional fold appearing on the lateral side, but collapsing with the pouch side fold before midline. HH and lateral folds have all collapsed into one fold in the mutant. vi) Schematics displaying the collapse of HH and HP folds and the LF. E) Simulation in (B), with 50 per cent growth reduction on hinge, top view ay 84 hr AEL, lateral cross-section point marked. ii) Lateral cross section. Red arrowhead marks the NH fold, black arrowheads mark the laterally initiating folds, reminiscent of HH, HP folds, or the LF. Scale bar on ii valid for both, 50 micrometres. F) Simulation results for reduced hinge growth at i) 25% of wild type, ii) 75 % of wild type. All other simulation parameters same as Fig. 5D. Panel structure, colour coding and scale bars are same as (B). G) *wingless* mutant phenotypes with emergence of a residual small peak within the grove of the ventral side fold of the mutant, highlighting one fold is lost as a result of HH and HP folds merging. NH, HH and HP folds are marked in red, green and blue arrowheads, respectively. See also Supplementary Figure 6.

## Discussion

Here we present a computational model of tissue growth and morphogenesis, incorporating spatial and temporal heterogeneity in growth rates and orientations, basement membrane (BM) mechanics and remodelling, and physical property heterogeneities within different layers of the tissue. Coupled with the stiff apical surface and BM mechanics, we demonstrate that the planar differential growth rates are key in defining the positions of epithelial folds of the wing disc. Upon identifying early growth phases as sufficient for initiation of fold morphology, we make predictions on the emergent morphology upon perturbation of early growth rates. By changing only the planar differential growth rates *in silico*, we successfully predict the morphology of a *wingless* mutant *in vivo*. With our computational analysis, we propose a novel mechanism whereby planar differential growth rates define epithelial fold initiation positions.

Growth under confinement can induce buckling, a uniform system bringing about uniform symmetrical ripples (Karzbrun et al., 2018; Pocivavsek et al., 2008; Shyer et al., 2013; Wang and Zhao, 2015). Heterogeneities in the physical properties of different layers within a uniformly growing tissue can alter the emerging fold patterns, and therefore be a viable mechanism for tissue patterning. Such symmetry breaking requires external resistance, and non-symmetric initial tissue shape (Fig. 3A, S2C). Still, in the wing disc, uniform growth is not sufficient to generate the experimentally observed fold morphology. Investigating the growth heterogeneities, we show that there is differential growth in the plane of the wing disc, and this can further alter the emerging fold morphology (Fig 4). When combined with structural requirements on BM, apical resistance, and the tissue stiffness, our model predicts that planar differential growth rates determine the fold initiation positions (Fig. 5D).

We show the structural requirement on the BM is that a stiff, elastic BM is necessary for the emergence of the correct fold pattern. Once this elastic BM is defined sufficiently stiffer than the cellular layer (8 times the apical stiffness and above), the correct pattern can be obtained with a range of parameters (Fig. S6A). The wing disc BM has high spatial and temporal variability (Fig. 5A). Our prediction that a large range of BM parameter space would induce correct fold pattern reflects that the wing disc would maintain robust fold patterns against the noise in BM structure.

Our simulations show that some form of viscous resistance on apical surface is necessary to facilitate buckling. The apical ECM has a different composition (Ray et al., 2015) and likely independent dynamics from the BM. Previous modelling work has already shown elastic apical ECM behaviour does not generate experimentally observed wing disc responses upon stretch (Keller et al., 2018). The composition of the contents of the lumen, and any possible tethering interactions with the peripodial layer (Gibson and Schubiger, 2000), would resist apical surface movement with fluid like mechanics. These interactions could provide the necessary viscous resistance, therefore be functional in tissue folding.

The structural requirement on tissue stiffness profile is that the apical layer should be stiffer than the rest of the cell body. The stiff apical layer, with consequent faster and larger emergent growth under constraints, brings about a differential growth along the apical-basal axis of the tissue. This difference then facilitates emergence of sharp apical indentations, in a similar mechanism to the emergence of brain cortex sulci (Tallinen et al., 2014). Similar to other organs (Tallinen et al., 2016), the tissue shape is also influential, as the folds do not emerge at correct morphology on a round tissue (Fig. S5E).

Upon fulfilling certain structural pre-requisites, our model predicts that planar differential growth rates determine the initiation of precise fold pattern on the complex wing disc epithelia. The ultimate validation for mechanisms suggested by a computational model is to challenge the model to predict emergent behaviour of perturbations. One such perturbation, *spade*^*flag*^ mutant of the *wingless* gene (Neumann and Cohen, 1996; Rodríguez Dd et al., 2002), has reduced growth on the wing hinge. All other conditions being equal, by changing only the planar growth rates *in silico*, we predict the mutant morphology will present two folds instead of three. Later, we validate this prediction *ex-vivo* (Fig. 6B-D). This emphasises the importance of planar differential growth in defining fold morphology. The phenotype is dose dependent and the correct morphology attempts to emerge from lateral regions in the mutant simulations, indicating that there is a minimum growth requirement, below which there simply is not enough material to support three folds in the tissue.

While our model is able to successfully generate the three-fold pattern, the timing of fold initiation in our simulations deviate somewhat from the experimental observations (65 hours AEL in simulations compared to 76-80 hours AEL (Sui et al., 2018) *in vivo*, Fig. S4E). Temporal dynamics of tissue and BM physical properties could lead to the offset in our fold initiation timing. BM demonstrates complex structural variability through the development time frame of our interest (Fig. 5A), and it is not possible to assume monotonically increasing or decreasing stiffness of the BM, without extensive further studies. The stiffness of the tissue itself is most likely altered through the same period, given that the residual tension of the tissue has been shown to reduce over time (Rauskolb et al., 2014). The fact that we can observe fold initiation in the simulations prior to emergence of the folds in the experiments indicates the accumulation of cell mass due to planar differential growth is sufficient for fold initiation, yet the tissue may not have reached the enabling physical state before 76 hours AEL.

At later stages of the simulations, the successfully initiated hinge folds do not progress into fully established folds (Fig. S5H), and the simulations deviate from the measured contour length of the experiments (Fig. S5B). Our simulations suggest that once the folds are initiated with planar differential growth rates, additional mechanisms should be activated to progress these indentations into folds. Indeed, cell shortening, alteration of the microtubule and actin networks, modification of both interaction with the BM through integrins and the BM structure itself through MMPs, have all been reported as necessary requirements for progression of wing disc folding (Shen et al., 2008; Sui et al., 2012, 2018). Our findings set the scene for further theoretical and experimental investigation on the feedback mechanisms between the morphology of the apical surface, and the signalling pathways regulating local growth, cell shape and BM interactions. In our model, we impose the growth patterns from experimental observation. Further analysis is needed in order to determine the coupling between this growth distribution and other chemical, genetic or chemical factors.

Our results suggest that planar differential growth is a novel mechanism for determining tissue fold positions, independent of active force generation. With wider implications, we suggest that the growth patterns giving tissues their final size can also regulate their architecture. We show that forces may not always result in instantaneous morphology changes, but can result in delayed morphogenesis. Stresses may accumulate early during development, even without any obvious changes in tissue morphology, but these may be critical for the precise sculpting of the tissue later in development.

## Methods

### *Drosophila* strains

For clone generation, tissue shape and morphology measurements: hsflp;; (BDSC8862) and w;; *Act*<CD2<GAL4, UAS-GFP “GFP on 3 Flipout (Neufeld et al., 1998). For wildtype height measurements: *yellow white* (*yw*;;) (BDSC). For *wingless* mutant analysis:; *wingless*^*spd-fg*^; (BDSC1005).

### Clone generation for growth analysis

To generate heat shock flip-out GFP clones of the correct density for growth rate analysis, the following regimes were used: for growth rates at 48–72h, 56-80h and 64–88h, heat shock was performed at 48h, 54h or 64h AEL respectively, for 12-20 min and dissected 24h later. For growth rates at 72–96h, heat shock was performed at 72h for 10 min and wing discs dissected 24h later. All heat shocks were carried out at 37°C

### Immunostaining and imaging of wing imaginal discs

Larval wing imaginal discs were dissected and stained as per the procedure described in Gaul et al., 1992. In brief, wing discs were dissected at the appropriate age in ice cold PBS for up to 15 minutes and fixed in 4% formaldehyde in PBS, at room temperature, for 30 minutes.

For *wingless* mutants, fixed discs were repeatedly washed within a 40 minute period in 0.3% PBT, followed by repeated washes with 0.5% BSA, 0.3% PBT for a further 40 minutes. Primary antibody, mouse anti-Wingless was prepared in 0.5% BSA, 0.3% PBT at 1:100 concentration and incubated overnight at 4°C. Washes were repeated as prior to primary antibody incubation. Secondary antibody, goat anti-mouse RRX (JacksonImmunoResearch) (1:500), Alexa fluor 647-Phalloidin (Cell Signalling and Life Technologies) (1:20) and Hoechst (Sigma-Aldrich) (1:500) were prepared in 0.5% BSA, 0.3% PBT and incubated for 1 hour at room temperature. Wing discs were washed repeatedly for 1 hour in 0.3% PBT, prior to rinsing in PBS.

For *yw* larva used for height measurements and flip-out clone larva used for growth rate analysis, dissected and fixed wing discs were washed in 0.3% PBT repetitively for 20 minutes, then immediately incubated with Alexa fluor 647-Phalloidin (Cell Signalling and Life Technologies) (1:20) and Hoechst (Sigma-Aldrich) (1:500) in 0.3% PBT for 15 minutes at room temperature. Wing discs were washed repetitively for 30-40 minutes, and then rinsed with PBS.

Fixed and stained wing discs were mounted in fluoromount G Slide mounting medium (SouthernBiotech) for imaging.

Wing discs were imaged on a Leica SP5 and SP8 inverted confocal microscope with a 40X oil objective at 1-2X zoom, 0.341 µm depth resolution and 512 by 512 or 1024 by 1024 pixel resolution.

### Electron microscopy

Wing discs were fixed in 2% formaldehyde/ 1.5% glutaraldehyde in PBS for 30 minutes prior to being flat, sandwich-embedded in 2.8% low melting point agarose dissolved in PBS. Once set, asymmetric cubes of agarose were cut out containing the wing discs and they were secondarily fixed for 1 hour in 1% osmium tetroxide/1.5% potassium ferricyanide at 4°C. Further fixation and contrast enhancement was achieved with, 1% thiocarbohydrazide for 20 minutes, 2% osmium tetroxide for 30 minutes, 1% uranyl acetate overnight at 4°C and lead aspartate for 30 minutes at 60°C, with extensive washes in double distilled water between incubations. Samples were then dehydrated in increasing concentration of ethanol solutions and embedded in Epon resin. The 70nm ultrathin resin sections were cut with a diamond knife (Diatome) using an ultramicrotome (UC7; Leica) and sections were collected on formvar-coated slot grids. Discs were imaged using a 120kV transmission electron microscope (Tecnai T12; FEI) equipped with a ccd camera (Morada; Olympus SIS).

### Tissue dimension measurements

The tissue size is measured from maximum projection images. DV length is defined as the longest axis from ventral tip of the pouch to the dorsal tip of the notum. AP length is measured to be the longest axis of the tissue perpendicular to the measured DV axis. The number of discs measured for each stage are: 4 discs at 48 hours AEL; 32 discs for early stages with no fold initiation; 38 discs for DV and 32 discs for AP for middle stages with some fold initiation; 22 discs for DV contour length, 31 discs for DV length and 17 discs for AP length for 96 hours AEL discs. Fold positions are measured at the longest axis in tissue midline, corresponding to the axis of DV length measurement. Each fold position is normalised to DV length, dorsal tip being 0 and ventral tip 1. The NH fold position is averaged from 19 wing discs, HH fold from 26 and HP fold from 16 discs.

### Growth rate analysis

The clone position, aspect ratio and orientation were calculated with automated segmentation and ellipse fitting, number of nuclei in each clone was counted manually, all using ImageJ. To convert the growth information of each clone to spatio-temporal maps of tissue growth, we go through age classification of the wing disc, alignment of wing morphology to average, binning the clone positions on a 2D projection of the tissue, and averaging data points, followed by overlaying the pouch growth rates from the literature(Mao et al., 2013), to generate the growth maps of Figure 4B. Next, the data points on the growth maps are intra/extrapolated to cover the empty spaces of the map grid, to allow for the maps to be smoothly read during the simulations (Fig. S3D-E).

The wing discs are divided into three age groups depending on their morphology, wing discs with no visible fold initiation, except for minor actin accumulation on the fold region, corresponding approximately to up to 80 hours AEL age (Fig. S3Ai); wing discs with at least one, mostly two to three initiated folds, but not reached to fully folded morphology, corresponding to 80-88 hours AEL age (Fig. S3aii-iv); and finally, wing discs with all three folds formed, corresponding to 96 hours AEL age. The age definitions are not clear-cut at all times, therefore we will refer to disc growth periods as early, mid and late phase, referring to the above morphological characterisation (Fig. S3A).

Upon division of the data points into age groups, where the wing disc has any markers for initiation of the first (HH) fold, the HH fold position of the individual disc is aligned to the average HH fold position, and the position of the clones updated (Fig. S3B). This alignment step is not applicable to wing-discs of the earliest growth phase, where no folds are visible. Next, all the clones for a selected time point are binned on a 20 by 10 grid, according to their normalised planar positions within the bounding box of the tissue (Fig. 4Aiii). Any grid bin with less than two data points is treated as empty. The growth data from the clones in each bin are averaged, Gaussian average is applied to number of nuclei, and the orientation angles, the orientation of the long axis of the fitted ellipse, are defined to be within π/2 degrees of each other before taking a Gaussian average. The aspect ratios are averaged with geometric average. Once the maps are generated, previously measured pouch growth rates(Mao et al., 2013) are aligned on top, defining the position of the HP fold as the dorsal tip of the pouch and using the pouch sizes measured in this study (Fig. S3C), and the maps presented in Figure 4B are obtained.

The simulation requires a complete map, without gaps, such that each element can read its growth rate at each time step. Therefore, we fill the empty points of the grid by interpolating the existing measurements. Starting from the centre of the grid and moving out radially (Fig. S3Di), once an empty grid point is detected, all the populated points within its eight immediate neighbours are averaged to fill the grid point (Fig. S3Dii). The order of filling is of significance, as once a point is filled with averaging the neighbours, it will be counted as a populated point in following iterations. This allows us to fill the grid points at all regions, and we obtain the maps in Figure S3E. Of note, the corner points are not necessarily sampled in the simulation, as the emergent simulation tissue shape is similar to that of the experiments, nevertheless, the map should cover a slightly larger area then the immediate experimental boundary to ensure continuity. For an example of the region sampled throughout the simulations, see Figure 6C. While reading the growth rates from these maps, each element of the simulation takes its centre point normalised to tissue bounding box, reads the closest four corner values from the growth and orientation maps, and interpolates the actual growth rate/orientation to apply depending on its distance from each of the four corners.

Growth in apical-basal axis is calculated from measurement of tissue thickness (Fig. 5Hii, S5F). The growth rate is directly calculated form the height increase, yet one complexity here became the pseudostratification of the tissue.

As the tissue grows, the nuclei become pseudostratified, first in the pouch, followed by the notum (Fig. 5Fiii-vii, S5J). Coinciding with the relative pouch thickness increase, the pseudostratification is visible in the pouch region as early as the initiation of the HH fold as an apical indentation (Fig. 5Fiv), while notum nuclei are still organised in a single layer (Fig. 5Fiii). As the development progresses, the pseudostratification can be seen everywhere, the extent being significantly higher in the pouch (Fig. 5Fvi-vii). The measurements demonstrate tissue thickness can increase without pseudostratification (Fig. 5Fiv), indicating addition of material to tissue height independent of cell division, i.e. the nuclei count from which we derive our planar growth rates. On the other hand, the pouch region of the tissue increases in height faster, to a greater extent, and pseudostratification is more predominant in this region (Fig. 5Fiv-vi), which should influence our definition of planar growth. To account for this difference, we allowed for tissue height increase to the level of the notum thickening, without altering the planar growth rates, reflecting the cell height increase independent of nuclei stratification. The additional height increase observed for the pouch region, the difference between the notum and pouch height increases, is reduced from the planar growth rates, “using up” the increase in nuclei numbers.

### The model definition

#### Elastic and viscous forces

The finite element model defines the tissue elastic properties as a neo-Hookean material, and the viscoelastic balance equations

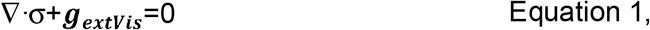

are solved on the whole tissue, with σ the Cauchy stress-tensor and *g*_*extVis*_ the external viscous forces. This viscous resistance is defined in terms of a viscous drag proportional to exposed surface area and displacement rate. These equations are discretised with finite elements and solved with an implicit integration scheme. A Newton-Raphson numerical procedure is employed, the emerging sparse matrix is solved using PARDISO solver (De Coninck et al., 2016; Kourounis et al., 2018; Verbosio et al., 2017). The tissue is assumed as a neo-Hookean material, where the second Piola-Kirshhoff stress tensor ****S**** is dependent on the strains as follows:

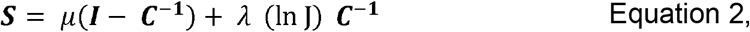

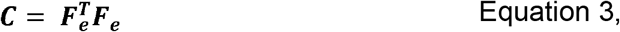

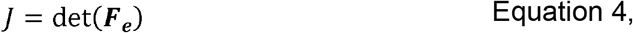

where ***C*** is the right Cauchy-Green deformation tensor, calculated from the elastic component of the deformation gradient, ***F***_***e***_, J is the determinant of ***F***_***e***_, and λ and μ are the Lamé constants defining the physical properties of the tissue. Cauchy stress tensor and the consequent elastic nodal force vector **g**_**e**_ are obtained following standard numerical elasticity procedures with shape function definitions (Bonet and Wood, 2008). Elastic and viscous forces are calculated within the implicit iterations, whereas evolution of the tissue in terms of growth, remodelling, and nodal interactions such as adhesion and binding, are calculated at the beginning of each time step, and kept constant during the iterations of the numerical solution procedure.

External viscous drag of a node *i* is calculated as a nodal drag force, proportional to the external viscous resistance of the environment and both velocity and exposed surface area of the node:

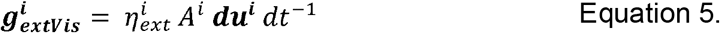

Here, 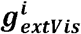, is the vector for external viscosity forces for node *i*, and the nodal external viscous forces are collated in system external viscous forces vector *g*_*extVis*_. Here, 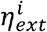 is the external viscous resistance coefficient for node *i*, *A*^*i*^ is the area contribution of node *i*, dt is the time step, and ***du***^*i*^ is the nodal displacement increment. For each element *e*, the elemental surface area is calculated and divided equally among the nodes constructing the surface. Area per node *A*^*i*^ is calculated as the sum of nodal surface areas of all elements owning the node *i*:

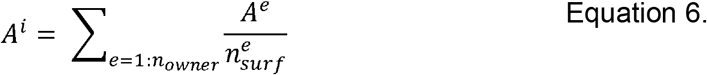

Here, n_owner_ is the number of elements connected to node *i*, *A*^*e*^ is the exposed surface area of interest of element *e*, and 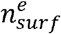 is the number of nodes that element *e* has on its exposed surface of interest (Fig. S2A). The area is calculated outside the numerical iteration, and based on the area of the previous time step.

#### Growth

Growth is incorporated into the simulations by resorting to a multiplicative decomposition of the deformation gradient as follows

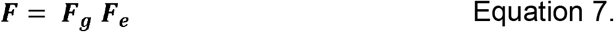

Here, ***F*** is the 3 by 3 deformation gradient matrix calculated from the reference and current coordinates of an element, ***F***_***g***_ is the growth deformation gradient representing the total growth of the element up to the current time step, and **F**_**e**_ becomes the residual elastic deformation gradient. The strain, corresponding stresses, and the resulting nodal forces, are calculated via ***F***_***e***_ (Fig. 2B). Note that ***F***_***g***_ is applied on the reference configuration of the element. The value of ***F***_***g***_ is updated at each time step *dt*, depending on the input growth rate, growth orientation, and the current rigid body rotations of the element following the equation:

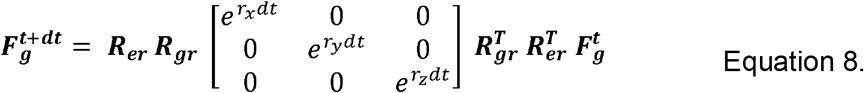

Here, *r*_*x,y,z*_ are the growth rates in the local coordinates of the element, *dt* is the time step, ***R***_***gr***_ is the rotation matrix for growth orientation as specified in the growth input, and ***R***_***er***_ is the rotation matrix associated with the current elemental rotation in the plane of the tissue. The growth orientations are calculated from the maximum projection of the experimental images on the xy-plane (see Methods: Growth analysis). The current rotation of the element around z-axis is corrected in order to ensure that the orientation of the growth follows the xy-plane of the tissue. At the beginning of the simulations, the local coordinate system of each element is aligned with the world coordinate system. During the simulation, the local coordinate system of the element could deviate from the world coordinates, due to rigid body rotations, imposed by the deformations of the surrounding tissue. Any rotation around the z-axis, changing the xy-plane of the element, should be accounted for, so that the element will continue growing on the desired orientation in the world coordinates. On the other hand, the tilt of the z-axis itself should be ignored; an element with tilted apical-basal (AB) axis should not start “elongating” in the AB direction (Fig. S2B). To obtain the rotation matrix ***R***_***er***_, first the current rigid body rotation of the element is calculated from the deformation gradient via single value decomposition such that

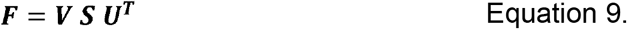

Then the rotation matrix corresponding to the rigid body rotation of the deformed element can be obtained from

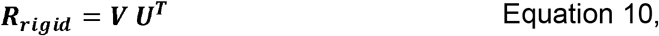

and the angle of rotation around the z axis is extracted from the calculated rigid body rotation matrix from

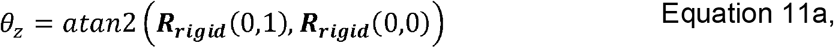

where

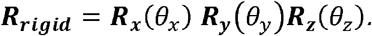

The elemental rotation matrix ***R***_***er***_ is then calculated for correcting the obtain rotation in z, as such, ***R***_***er***_ is rotation around z axis by −*θ*_*z*_.

#### Remodelling

While the cellular elements of the tissue grow with specified growth rates and orientations, the BM grows by remodelling. The application of remodelling follows the logic of equations 7-8, with the growth increment and related rotations defined from deformations, rather than an input growth profile. As such, the rigid body rotation correction, ***R***_***er***_, is not necessary for BM remodelling, and the equivalent of growth orientation (***R***_***gr***_) is obtained through the elastic deformation orientation as follows: The remodelling growth at each time step is obtained via eigen value decomposition of the Green strain matrix ***E*** of the element, and the deformations on the principal axes are calculated via

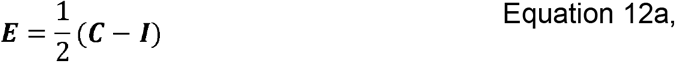

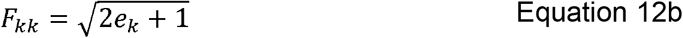

Here, *F*_*kk*_ is the current deformation along the principal axis k (such that a 50 per cent stretch will give 1.5), and *e*_*k*_ is the k^th^ eigen value of ***E*** matrix, where k =1:3 in 3D. The deformation after the relaxation to be observed within the current time step *dt* is then calculated from the given remodelling half-life (t_1/2_) as follows:

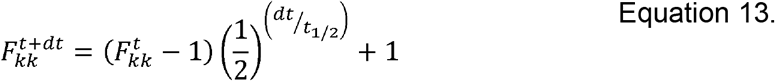

The calculated new deformation is then converted to a growth increment, with the orientation of the growth defined by the eigenvectors matrix **V**:

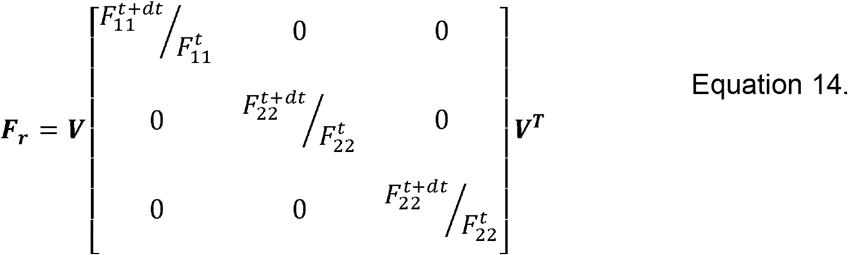

The remodelling serves to mimic BM remodelling carried out by the cells as the tissue grows, adding or removing material from the BM layer as needed. Therefore, remodelling of BM is done without volume conservation (determinant of ***F***_***r***_ can deviate from unity). Similar to Eq. 8, the remodelling growth increment ***F***_***r***_ is added to total growth deformation gradient ***F***_***g***_:

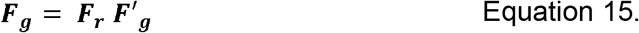

Moreover, as the BM is stretched and the new BM is allocated, there is no evidence that the BM should be getting thinner due to elastic forces and this thinning being reflected into the emergent shape, on the contrary, BM does get thicker with age. As such, remodelling is limited to plane of the tissue. BM is not remodelled in the apical-basal axis, only the x & y dimensions of the strain matrix are included in the decomposition, and the range of k in equations 9–11 are limited to k = 1:2 in 2D, excluding local z coordinates of the element, which is aligned with apical-basal axis.

#### Node-node interactions

Nodes can interact with each other via direct adhesions, or a packing hard-wall potential to ensure volume exclusion. The same threshold is used for detecting nodal adhesion or for applying the hard-wall potential. The packing forces are used mainly for cases where adhesion between the nodes is not feasible, such as interactions of nodes that share elements, where the adhesion would cause the shared element to flip.

The hard wall potential is defined to ensure volume exclusion as the elements move too close to each other. The potential is applied on a nodal basis. The threshold of repulsion force application, *τ*, is dynamic in the simulations, scaling to the average side length of an element in the vicinity of potential node-node interaction. A sigmoid function is selected to have a repulsion force with a continuous derivative, necessary for convergence in the implicit N-R numerical method

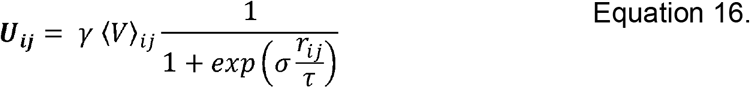

Here ***U***_***ij***_ is the force factor nodes *i* and *j*, γ is a scaling factor to convert the potential to forces relevant to the simulation forces, ⟨*V*⟩_*ij*_ is the average volume associated with nodes *i* and *j*, τ is the threshold distance for considering repulsion force, and *r*_*ij*_ is the distance between nodes *i* and *j* in 3D, and σ is a scale factor to define how much force will be applied at the threshold distance (Fig. S2C). A force equal to the potential but in opposing directions are applied on each node *i* and *j*. The threshold distance is defined to be 40 per cent of the average local side length, calculated on a 10 by 5 grid on the tissue xy bounding box. The parameters of the hard-wall potential are selected numerically in order to ensure volume exclusion, not necessitating a biological basis.

Additional to interacting with a hard wall potential, nodes within a close vicinity can bind to each other (Fig. S1D). Adhesion is defined by moving both nodes to the mid-point, and collapsing their degrees of freedom. The two nodes are assigned master and slave status arbitrarily. All the driving and drag forces of the slave node are carried on to the master node, and the Jacobian is updated accordingly. Upon obtaining the displacements, the displacement of the slave node is updated with the displacement of the master node (Eq. 17-20). This is equivalent to a master-slave treatment of the nodal constraint **x**_*slave*_=***x***_*master*_(Muñoz and Jelenić, 2004). One final check on the similar grounds is the adhesion of nodes on the same element, when two nodes come close to each other below a collapse threshold. This adhesion effectively collapses the degrees of freedom of the element, and protects the element from flipping. The threshold for the collapse is stricter than adhesion. It is defined on an elemental basis, and is defined as a distance below 10 per cent of the initial reference length between the two potentially collapsing nodes, as opposed to being proportional to the current average side length of the system (Fig. S2E). The residual vector ***g*** (sum of all nodal forces acting on the system) and Jacobian matrix ***K*** (derivatives of all nodal forces with respect to the nodal displacements) are modified as follows,

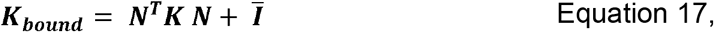

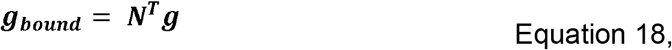

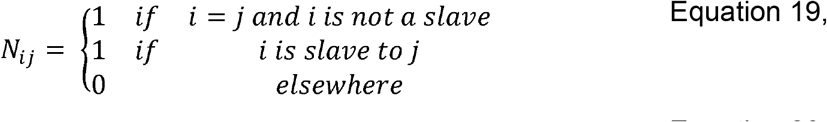

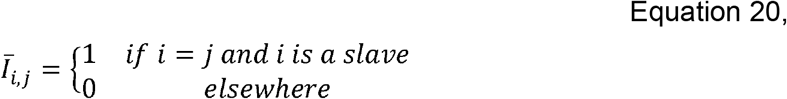

with i=1:3n_nodes_, j=1:3n_nodes_ looping over the degrees of freedom of the system. Upon calculation of displacements ***du*** with the new nodal forces, ***g***_***bound***_, and Jacobian, ***K***_***bound***_, the displacement of slave degrees of freedom are equated to their respective masters.

The same master slave definition is used for no-bending boundary condition. At the circumference, the basal node of each column of nodes is assigned as the master of all the remaining nodes of the column, and the degrees of freedom in x and y directions of slaves are fixed on the master (Fig. S2F).

#### Automated fold detection

The fold initiation is detected automatically by calculating the surface normal of all elements exposed on the apical or basal surface. If the surface normals within the vicinity of each other face in opposite directions (dot product being negative), the elements are defined to be on a folding curve. All the elements lying in between the two elements are also included on the fold surface (Fig. S2G). The threshold distance for identifying curved regions is selected as 3 micrometres. The threshold is selected to ensure detection of fold initiation that is clearly visible when the morphology is visualised, but do not assign fold initiation identity to elements at opposing sides of a possible curve peak. For the looser folds of the uniform growth rate simulations, a more generous threshold of 6 micrometres is used.

#### Simulation fold position scoring

Of the detected continuous fold initiations, for those that reach the midline, the positions are aligned with the experimental fold positions (Fig, 1Cii), such that the minimum deviation is calculated. If the simulation produces less than 3 folds, each missing fold is counted as 100% deviation. The score does not penalise for additional folds, such as those observed with uniform growth and explicit BM definition (Figure 5E). The deviation score also not check the fold morphology, such as the high hinge folds of uniform growth rates that reach taller than the pouch region (Fig. 3Ai). Therefore each low deviation value should be examined against both.

## Acknowledgements

We thank all the authors and Erik Sahai, Robert Tetley, Alejandra Guzman Herrera for revision of this manuscript. MT is funded by a Sir Henry Wellcome Fellowship (Grant No: 103095). MD is funded by a Marie Skłodowska-Curie Horizon 2020 Individual Fellowship (MRTGS). NJK is funded by a MRC PhD studentship. RB is funded as part of the CoMPLEX doctoral training program core studentship. JJB is supported by MRC funding to the MRC LMCB University Unit at UCL, award code MC_U12266B). JJM is funded by Spanish Ministry of Economy and Competitiveness (grant DPI2016-74929-R) and Generalitat de Catalunya (grant 2017 SGR 1278). YM is funded by a MRC Fellowship MR/L009056/1, a UCL Excellence Fellowship, and a NSFC International Young Scientist Fellowship 31650110472. This work was also supported by MRC funding to the MRC LMCB University Unit at UCL, award code MC_U12266B

