## Supplementary figures and images for "Planar differential growth rates determine the position of folds in complex epithelia"

### Supplementary Figure 1

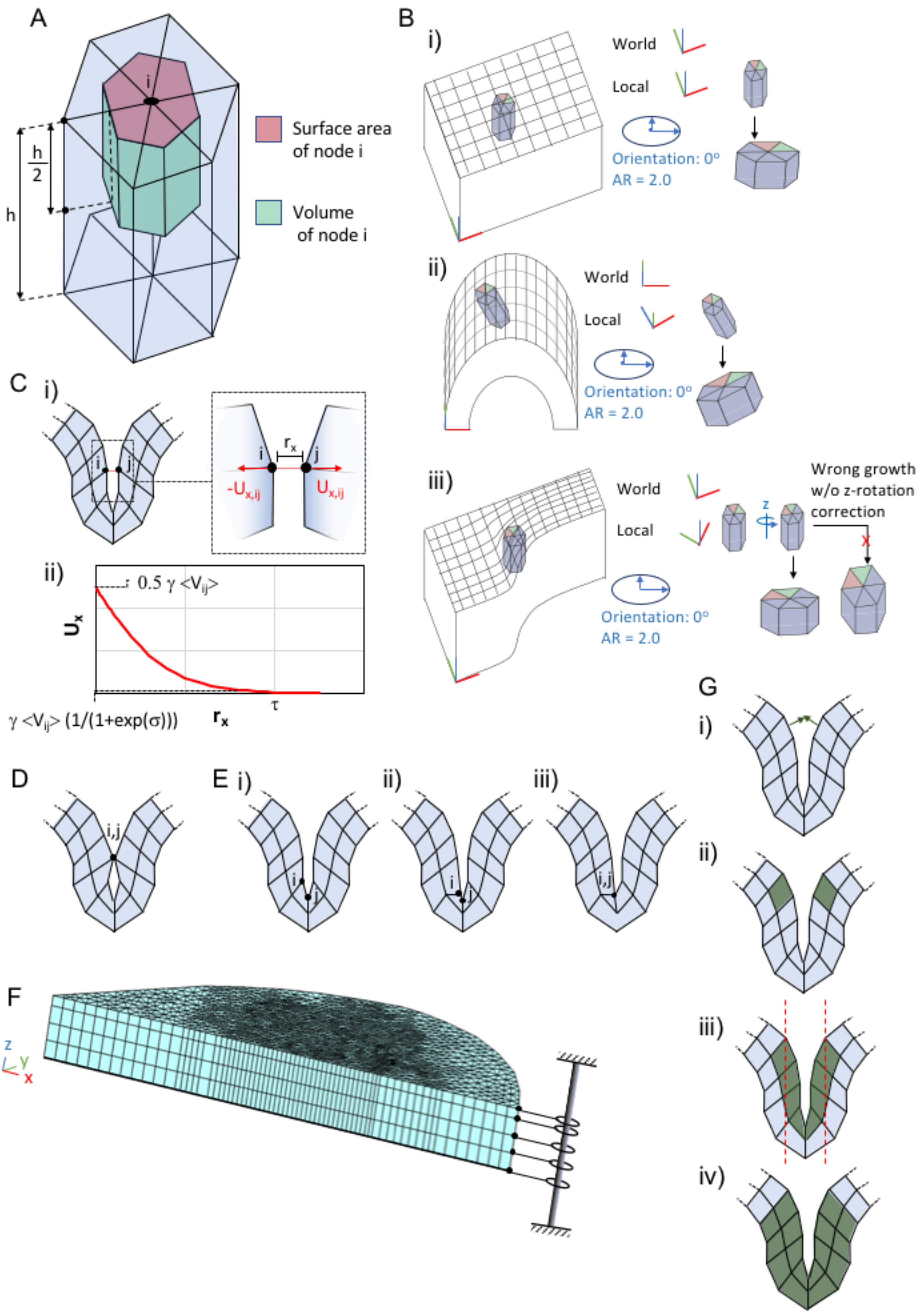

### Supplementary Figure 2

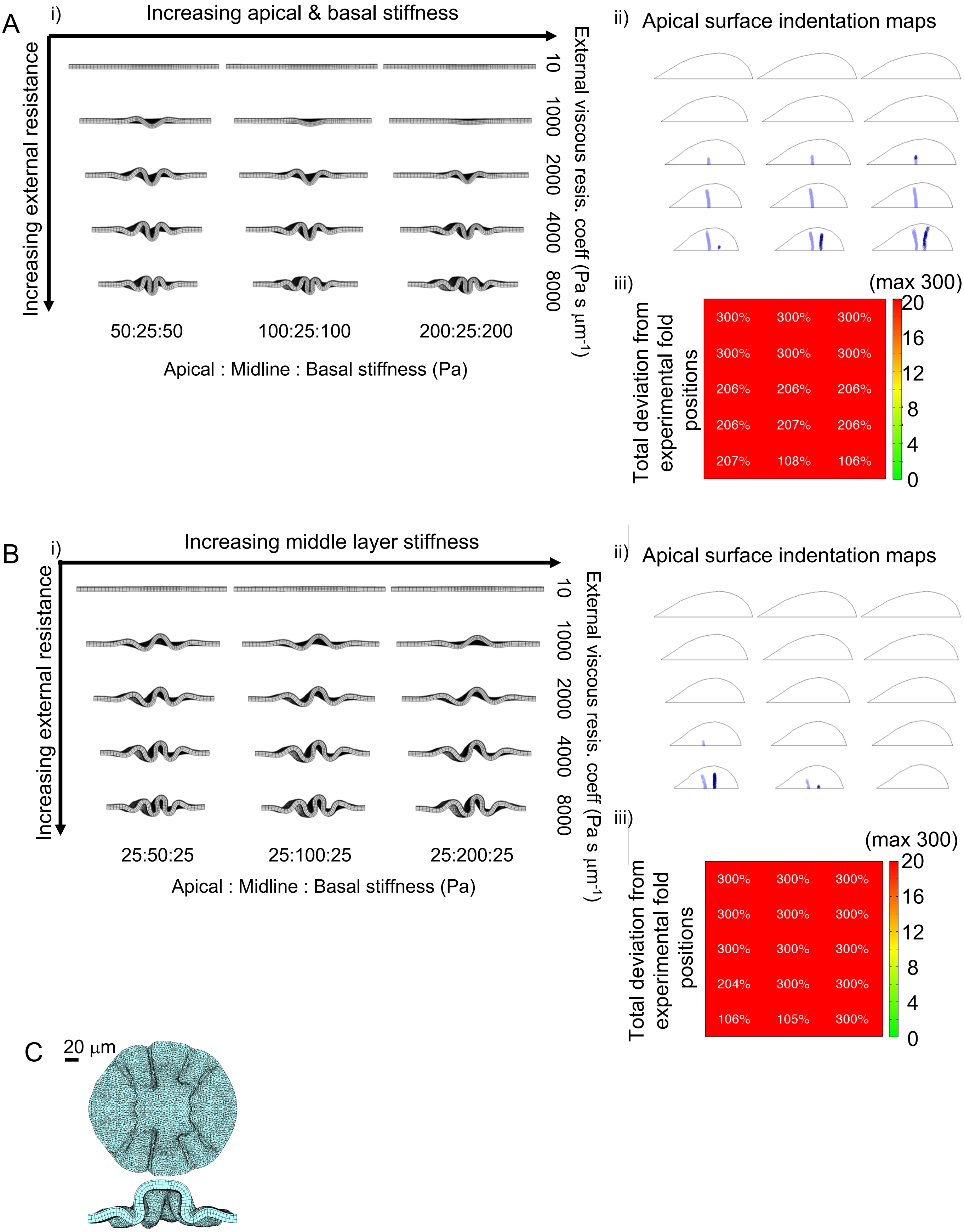

### Supplementary Figure 3

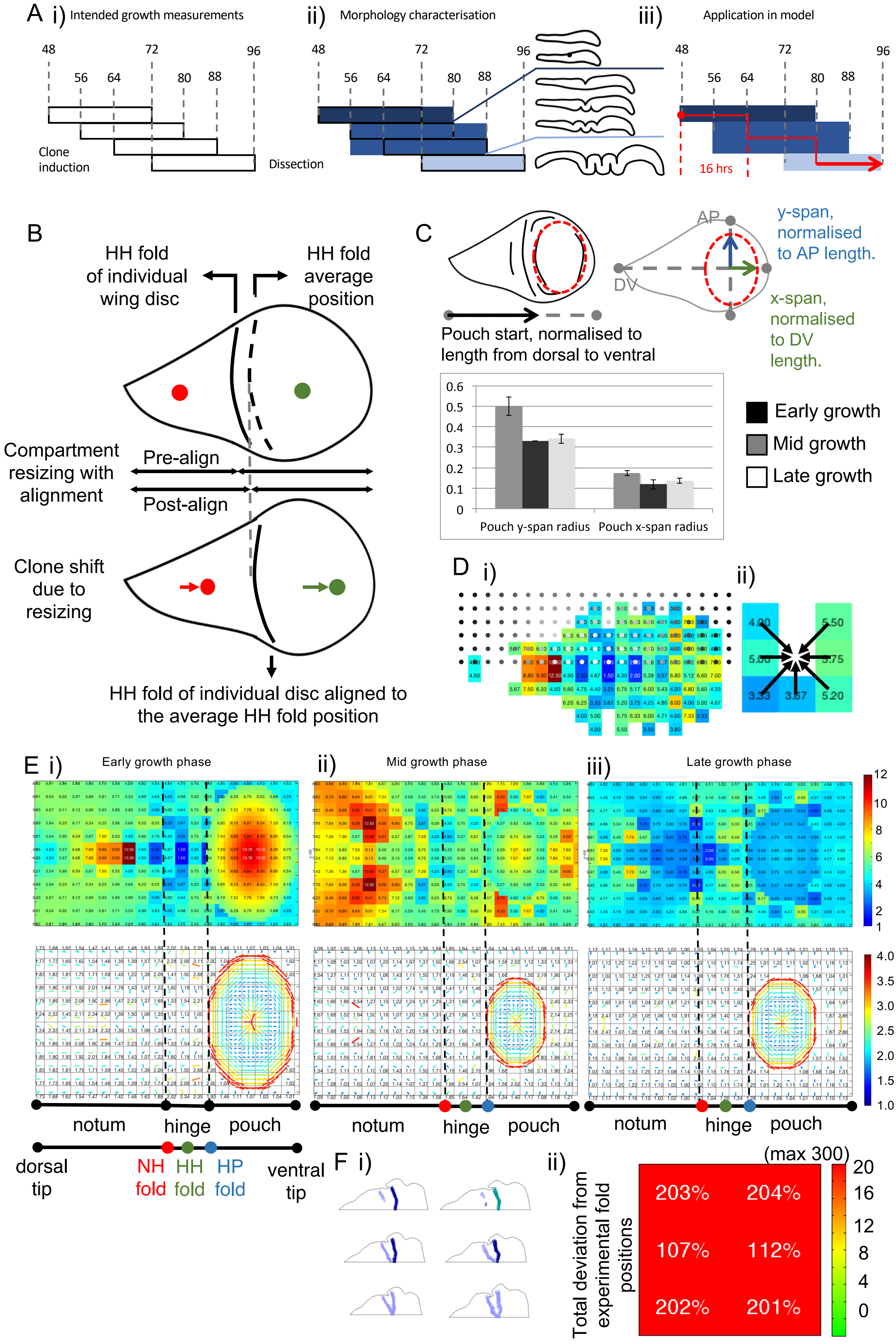

### Supplementary Figure 4

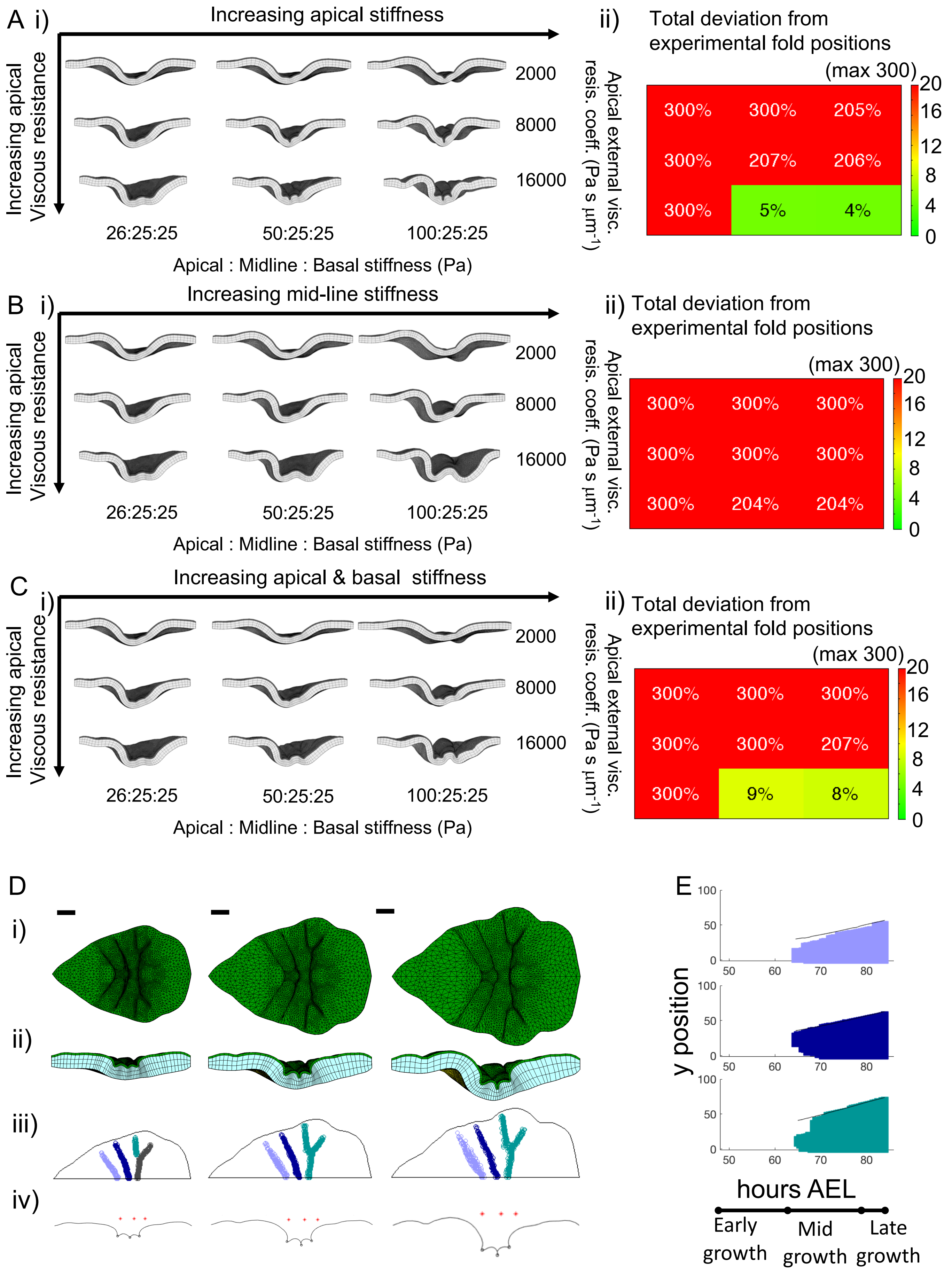

### Supplementary Figure 5

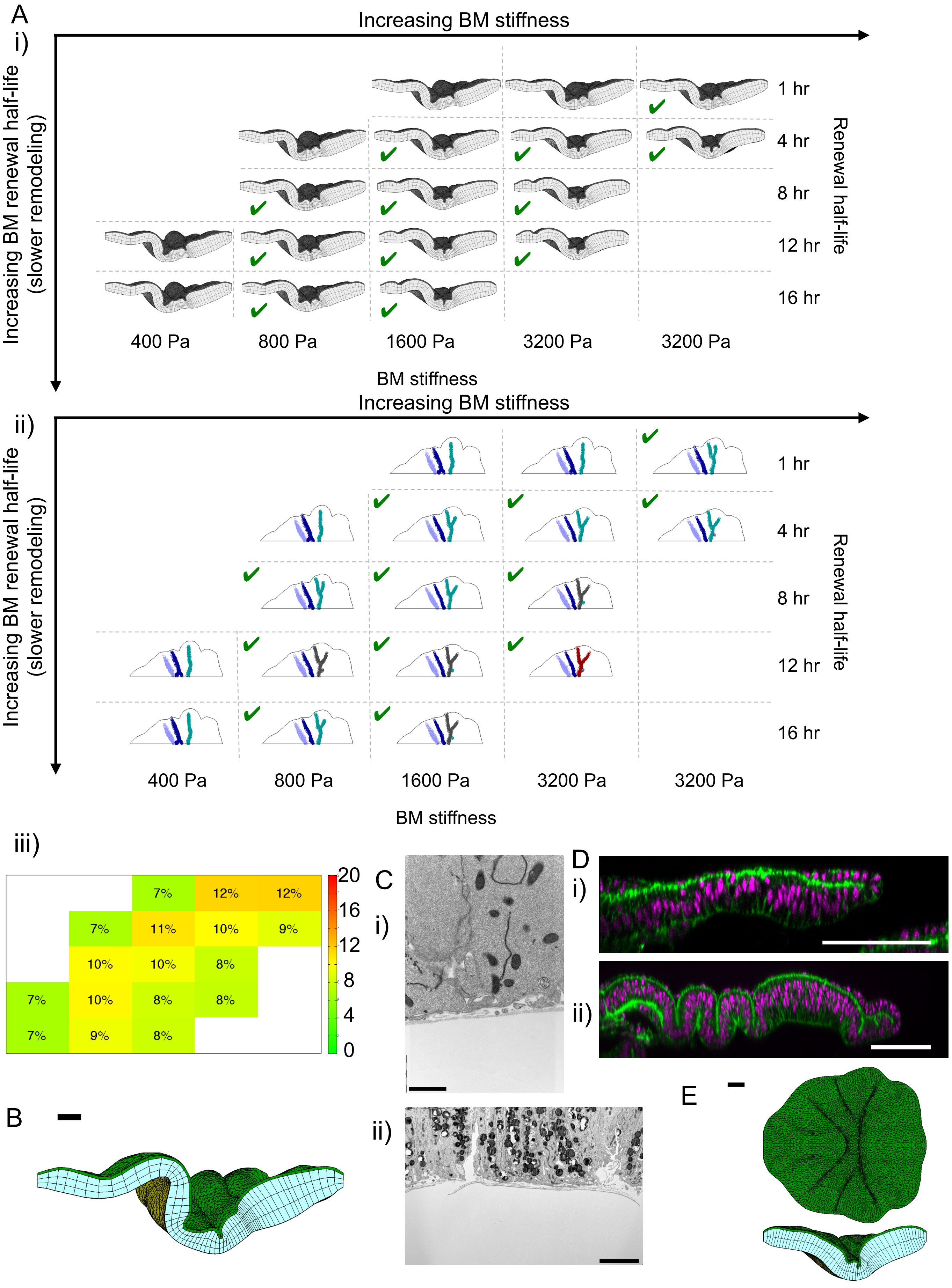

### Supplementary Figure 6

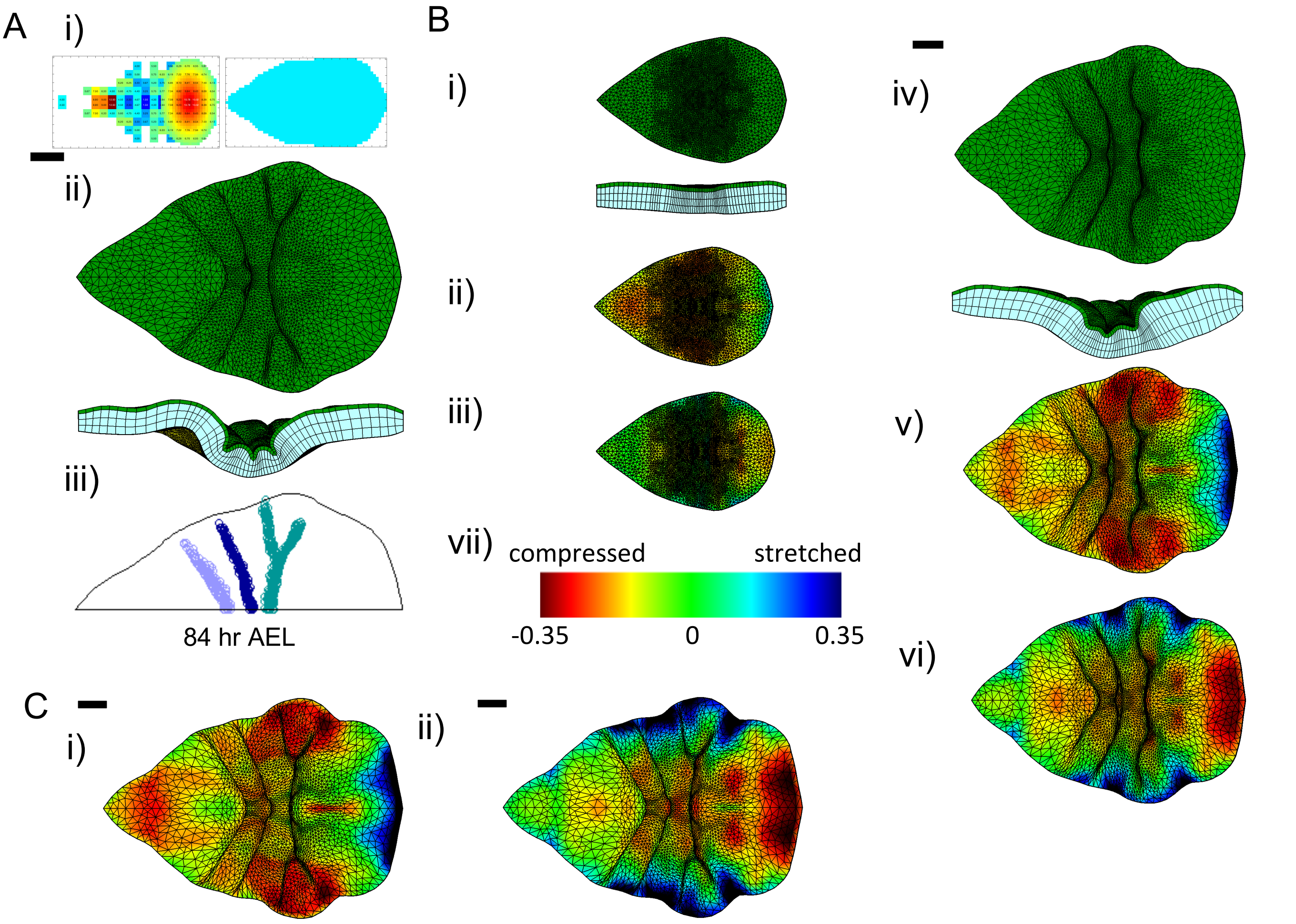
