## Supplementary Information for "Planar differential growth rates determine the position of folds in complex epithelia"

**Supplementary Figure Legends**

**Supplementary Figure 1.** Related to figure 2. Modelling method details. A) Schematic for the externally exposed surface area and volume allocation to nodes. Node i is shared among six triangular prism elements, the total surface area highlighted in red, volume on green. B) Schematics for the oriented growth and related rotational corrections. A growth of doubling in volume, parallel to x-axis, with aspect ratio of 2 is demonstrated for all cases. All coordinate axes display x in red, y in green and z in blue. In the simulations, x is aligned with the DV axis, y with AP axis and z with AB axis of the tissue. i) Simple scenario where the world and local coordinates are aligned. ii) The elements have a rigid body rotation that deviates the local z-axis from the world coordinates, x & y are aligned. The growth follows local coordinate system, and the rigid body rotation is not accounted for. iii) The case where due to deformation in the tissue, the elements have gone through a rigid body rotation around the z-axis. The growth orientation is corrected for the rotation around z, and growth on x-y plane is applied in world coordinates. The wrong emergent grown shape in the case when this rotation was ignored is shown for comparison. C) Schematic for the hard wall potential applied to ensure volume exclusion. i) packing forces between nodes i and j in x-axis. The dashed box is enlarged on the inset, the calculated potential is applied in the opposite directions on both nodes. ii) The packing potential with distance displayed, the parameters defining the potential function are marked. D) Schematic displaying the adhesion of nodes i and j, the initial configuration same as Ci, the nodes are carried to the mid-point and their degrees of freedom bound. E) Schematic for node collapse on elements with nodes approaching within a small threshold distance of each other, implemented to limit element flipping. i) Node configuration outside collapse limit, ii) nodes moved within the collapse limit of each other due to viscoelastic system forces. iii) Configuration after node collapse. In D and E, the schematics are for demonstration purposes only and distances are not to scale. F) A sample simulated initial mesh, displaying the symmetricity assumption and showing the simulated half. Schematic added on the simulation mesh to demonstrate the no-bending boundary condition at circumference. G) Schematic demonstrating the algorithm to detect fold initiation with element surface normals. The detection is carried out on elements with exposed surfaces on either apical or basal surfaces, apical surface is utilised in demonstration. i) two normals on elements (green arrows) are within the vicinity of each other and the angle between the normals is wider than the selected threshold. ii) Two elements are assigned to be on fold initiation regions. iii) All elements that have their apical surfaces within the bounding box of the identified element couple are marked to be on fold initiation surface. For fold identification on the basal surface, the bounding box will check for basal surfaces of the remaining elements. iv) The fold initiation region is extended to cover the whole tissue thickness.

**Supplementary Figure 2:** Related to figure 3. Alternative relative stiffness states of the tissue. Tissue morphology with, A) increasing relative stiffness of apical and basal surfaces and external viscous resistance; B) increasing relative stiffness of tissue midline, and external viscous resistance. Each panel demonstrates simulation results for a tissue growing from 48 hour AEL to 96 hours AEL, with uniform in-plane growth rates, as stated in Figure 3A. Images are taken the cross-section of the tissue midline at 96 hours AEL, ventral tip on the right. Row and Column organization same as Figure 3A. Both in A and B, ii) Apical indentation maps automatically identified from the curvature of facing surfaces, each continuous folding region is marked in a single colour. iii) Fold position deviations, calculated as sum of percentage deviation from each experimental fold at the tissue centre. Both i & ii calculated at same time points of (i), row column organisation is same as in (i). C) Simulation with same parameters as Figure 3B, on a symmetric, circular initial mesh. Snapshot is from 96 hours AEL, sagittal view as central cross-section, and dorsal tip to the right. Scale bar 20 micrometres.

**Supplementary Figure 3:** Related to figure 4. Details of the growth rate analysis methodology. A) Definition of the growth phases. i) The targeted experimental timings. ii) The morphological staging of the dissected wing discs according to the folding stage. iii) The timing for the application of the growth phases in the simulations. B) Schematic representing the alignment of the growth rate measurements for a given time period. HH fold position (dashed line) is calculated as the average of all the wing-discs utilized in growth rate measurements for the selected time period. The HH position of each individual wing disc (solid line) is aligned to the average position. The two sides of the fold are rescaled to fit within the average size of the tissue, moving the data points (red on notum side, and green on pouch side) in the process. C) Measurements for the size of the pouch at different time periods within 48 to 96 hours AEL. Top schematics demonstrate how the pouch position and size are normalised. D) The order in which the grid points are checked, and filled as necessary. i) Order goes from light to dark shades of dots on grid points. ii) Once an empty grid point (with no experimental data points) is reached, the existing data in immediate neighbours are averaged to cover the empty point. The order of sampling for empty grid points is of significance as filled regions contribute to the filling of their neighbours, thus enabling us to fill the extended patches of empty regions on the grid. E) The extended version of growth maps demonstrated in Figure 4B. Colour bars on the right hand side of the panes are valid for all. The measured fold positions, and the corresponding tissue compartments are marked on the heatmaps, the positions are measured for 72-88 hours AEL in i&ii, for 96 hours AEL in iii. F) i) Apical indentation maps and ii) fold position deviations of simulations with experimental growth rate, as presented in Figure 4C.

**Supplementary Figure 4:** Related to figure 5. A-C) i) The effects of apical viscous resistance coefficient and tissue stiffness heterogeneity on emergent morphology. The tissue has explicit BM definition at 1600 Pa stiffness and renewal half-life of 8 hour, basal viscous resistance coefficient is 10 Pa s μm^-1^. ii) Fold position deviations, calculated as sum of percentage deviation from each experimental fold at the tissue centre, row column organisation is same as in (i). A) Apical surface is stiffer than the rest of the cell body, columns: increasing apical viscous resistance coefficient, rows: increasing relative stiffness of apical surface. Simulation in lower right corner is detailed in Figure 5D. B) Simulations with tissue midline stiffer than the rest of the tissue, grid organisation same as (A). C) Simulations with tissue apical and basal surfaces stiffer than the rest of the cell body, grid organisation same as (A). D) Timeline of z-growth added simulation in Figure 5G and Movie 5. Snapshots are from 72, 78, and 84 hours AEL, respectively. Scale bar is 20 micrometres. i) top view, ii) sagittal view, iii) apical indentation maps. iv) The positions of the folds on the tissue cross section apical surface profile, the red stars mark the experimental fold positions measured for 72-88 hours AEL (Fig. 1Ci). E) Kymographs of apical indentations in time. Y position of all nodes falling into the three major folds at 84 hours AEL are plotted in time. Colour coding same as (Diii) at 84 hours AEL.

**Supplementary Figure 5:** Related to figure 5. A) The effects of BM stiffness and BM renewal half-life on emergent morphology. Rows, increasing BM stiffness, columns increasing renewal half-life (slower remodelling). i) Sagittal views displayed for each parameter combination at 84 hr AEL. ii) apical indentation maps. Acceptable parameter combinations marked with a green tick in (i&ii). Failing simulations form a forked HH fold, merging with the NH fold at tissue midline. iii) Percentage deviation of fold positions from the experimental positions measured at tissue midline at 36 hr AEL. B) The late stages of the simulation in Figure 5G, at 96 hours AEL. The initiated folds do not successfully progress into a fully folded morphology. The tissue goes through large scale buckling, the HH fold is opened up and hinge sinks well below the notum in z. Scale bar, 20 micrometres. C) The larger field view images of EM images (Figure 5A), i) 72 hr AEL, scale bar 1 micrometre, ii) 120 hr AEL, scale bar 5 micrometre. D) The larger field of view images where the pseudostratification images are taken from i) Figure 5Fiv-v, ii) Figure 5Fvi-vii. Scale bars 50 micrometres. E) Simulation on a circular initial tissue shape, with parameters same as Figure 5G. Scale bar 20 micrometres.

**Supplementary Figure 6:** Related to figure 6. The differential growth in early growth phases and related force accumulation is necessary and sufficient for correct morphology in wild type and mutant wing discs. A) Early growth rates applied for the initial 16 hours (48 to 64 hours AEL) of simulation, continued by uniform growth. i) The growth maps applied, colour coding same as Figure 4B. ii) Snapshots from 84 hours AEL, top and sagittal view, scale bar 20 micrometres. iii) Apical indentations map at tissue at 84 hours AEL, scale same as (ii). Simulation physical parameters are same as 5D. B) Simulation with experimental growth rates, with all accumulated forces relaxed at 58 hours AEL (10 hours into simulation). i) Tissue morphology at 58 hours AEL, immediately prior to relaxation of forces. ii) Strains accumulated in DV orientation, and iii) strains on AP orientation, colour coding in (vii). iv-vi) Simulation at 84 hours AEL, panel structure same as i-iii. C) Accumulated strains in wild type simulation, in Fig. 5H. i) DV stains, ii) AP strains, colour scale same as in (Bvii).

**Supplementary Movie Legends**

**Movie 1:** Uniform planar growth rates are not sufficient to generate experimental fold morphology, related to Figure 3: Tissue growing from 48 hour AEL to 96 hours AEL, with uniform in-plane growth rates. The growth rate is 0.033 hr^-1^ in DV, and 0.028 hr^-1^ in AP, as calculated from Figure 1C. Apical stiffness is 200 Pa (green), and stiffness for the rest of the cell body is 25 Pa (blue). Apical ECM/BM effects modelled as viscous resistances and external viscous resistance coefficient applied to both surfaces at 8000 Pa s μm^-1^. Scale bar is 20 micrometres. Simulation time depicted on the frames.

**Movie 2:** Emergent fold morphology with planar differential growth rates and a simplistic definition of BM, related to Figure 4: Tissue growing from 48 hour AEL to 84 hours AEL, with experimental planar growth rates (Fig. 4B). Apical stiffness is 200 Pa (green), and stiffness for the rest of the cell body is 25 Pa (blue). Apical ECM/BM effects modelled as viscous resistances and external viscous resistance coefficient applied to both surfaces at 8000 Pa s μm^-1^. Scale bar is 20 micrometres. Simulation time depicted on the frames.

**Movie 3:** Correct fold morphology emerges with planar differential growth rates and an explicit definition of the elastic BM, related to Figure 5(D): Tissue growing from 48 hour AEL to 84 hours AEL, with experimental planar growth rates (Fig. 4B) and an explicitly defined BM (yellow). Apical stiffness is 100 Pa (green), and stiffness for the rest of the cell body is 25 Pa (blue). Apical ECM effect modelled as a viscous resistance with external viscous resistance coefficient of 16000 Pa s μm^-1^. BM stiffness is 1600 Pa, and BM renewal half-life is 8 hours. Scale bar is 20 micrometres. Simulation time depicted on the frames.

**Movie 4:** Differential thickness increase confines the folds to the hinge region, related to Figure 5(G): Tissue growing from 48 hour AEL to 84 hours AEL, with experimental planar growth rates (Fig. 4B), differential tissue thickness increase (Fig. 5Fii, see methods), and an explicitly defined BM (yellow). Apical stiffness is 100 Pa (green), and stiffness for the rest of the cell body is 25 Pa (blue). Apical ECM effect modelled as a viscous resistance with external viscous resistance coefficient of 16000 Pa s m^-1^. BM stiffness is 1600 Pa, and BM renewal half-life is 8 hours. Scale bar is 20 micrometres. Simulation time depicted on the frames.

**Movie 5:** Predictions of the emergent morphology for *spd^fg^* mutation of the *wingless*, related to Figure 6: Tissue growing from 48 hour AEL to 84 hours AEL, with planar differential growth rates mimicking the *spd^fg^* mutation of the *wingless*. The growth of the hinge is reduced by 50 per cent for this particular simulation (Fig. 6A). There is differential tissue thickness increase (Fig. 5Fii, see methods). Explicitly defined BM (yellow) has stiffness of 1600 Pa, and renewal half-life of 8 hours. Apical stiffness is 100 Pa (green), and stiffness for the rest of the cell body is 25 Pa (blue). Apical ECM effect modelled as a viscous resistance with external viscous resistance coefficient of 16000 Pa s μm^-1^. Scale bar is 20 micrometres. Simulation time depicted on the frames.
